# Artesunate interacts with Vitamin D receptor to reverse mouse model of sepsis-induced immunosuppression via enhancing autophagy

**DOI:** 10.1101/2020.02.26.966143

**Authors:** Shenglan Shang, Jiaqi Wu, Xiaoli Li, Xin Liu, Pan Li, Chunli Zheng, Yonghua Wang, Songqing Liu, Jiang Zheng, Hong Zhou

## Abstract

**Background and Purpose:** Immunosuppression is the predominant cause of mortality for sepsis due to failure to eradicate invading pathogens. Unfortunately, no effective and specific drugs capable of reversing immunosuppression are available for clinical use. Increasing evidence implicates vitamin D receptor (VDR) involved in sepsis-induced immunosuppression. Herein, artesunate (AS) was discovered to reverse sepsis-induced immunosuppression and its molecular mechanism is investigated.

**Experimental Approach:** Effect of artesunate on sepsis-induced immunosuppression was investigated in mice and *in vitro*. VDR was predicted to be an interacted candidate of AS by bioinformatics predict, then identified using PCR and immunoblotting. VDR, *ATG16L1* and NF-κB p65 were modified to investigate the alteration of AS’s effect on pro-inflammatory cytokines release, bacteria clearance and autophagy activities in sepsis-induced immunosuppression.

**Key Results:** AS significantly reduced the mortality of cecal ligation and puncture (CLP)-induced sepsis immunosuppression mice challenged with *Pseudomonas Aeruginosa*, and enhanced proinflammatory cytokines release and bacterial clearance to reverse sepsis-induced immunosuppression *in vivo* and *in vitro*. Mechanically, AS interacted with VDR thereby inhibited the nuclear translocation of VDR, then influencing *ATG16L1* transcription and subsequent autophagy activity. In addition, AS inhibited physical interaction between VDR and NF-κB p65 in LPS tolerance macrophages, then promoted nuclear translocation of NF-κB p65, which activated the transcription of NF-κB p65 target genes such as pro-inflammatory cytokines.

**Conclusion and Implications:** Our findings provide an evidence that AS interacted with VDR to reverse sepsis-induced immunosuppression in an autophagy and NF-κB dependent way, highlighting a novel approach for sepsis treatment and drug repurposing of AS in the future.

## 1. Introduction

Sepsis is a leading cause of death worldwide (Angus & van der Poll, 2013); it arises when hosts response to pathogen infections such as bacterial, fungal, viral and parasitic infection and then injury their own tissues and organs (Hotchkiss, Monneret & Payen, 2013b). At present, severe acute respiratory syndrome coronavirus 2 (SARS-CoV-2) a newly recognized viruse, has spread rapidly throughout Wuhan (Hubei province) to other provinces in China and around the world, and has infected closed to 80,000 people in China and killed more than 2,000 untill Feb 26, 2020 because of no specific and effective drug.

In the course of sepsis, both innate and acquired immune systems are involved in its development, in which the mononuclear phagocyte system plays a crucial role. Although the timeline of immunological alterations in sepsis is not well understood, the suggestive evidence demonstrated that the development of sepsis involves two distinct stages, an initial cytokine storm (hyper-inflammation) and subsequent immunosuppression (hypo-inflammation), which exist concomitantly and dominate in different stage (Hotchkiss, Monneret & Payen, 2013a; Hutchins, Unsinger, Hotchkiss & Ayala, 2014). The uncontrolled cytokine storm is responsible for deaths occurring within the first few days, whereas immunosuppression predominantly causes mortality during the later stage of sepsis. In fact, more than 70% of patients die after the first three days of sepsis, which might due to the increased susceptibility to weakly virulent pathogens or opportunistic bacteria (e.g., *Pseudomonas aeruginosa*), indicating a failure of the host to eradicate invading pathogens (Hotchkiss, Monneret & Payen, 2013a; Skrupky, Kerby & Hotchkiss, 2011). For patients in immunosuppression stage, only less than 5% of monocytes/macrophages produce cytokines, which might be related to the disordered immune response in sepsis (Hotchkiss, Monneret & Payen, 2013a; Hutchins, Unsinger, Hotchkiss & Ayala, 2014), demonstrating the impaired function of monocytes/macrophages is tightly related to the decrease of removing bacteria. Considering the drugs based on the inhibition of critical pro-inflammatory mediators have failed, the therapeutic strategy to improve immunosuppression is the focus of sepsis treatment now (Cohen, 2002; Hotchkiss, Monneret & Payen, 2013a; Kox, Volk, Kox & Volk, 2000; Prucha, Zazula & Russwurm, 2017).

Autophagy is a fundamental cellular process involved in defense against infection by intracellular pathogens (Casanova, 2017). Although contradictory findings have been reported (Ho et al., 2016), accumulating evidence suggest that autophagy plays a key role in sepsis, and has been considered a potential target for sepsis-induced immunosuppression (Ren, Zhang, Wu & Yao, 2017). It is promising to focus on autophagy axis to unveil the mechanism of sepsis-induced immunosuppression and discover the novel immunoregulating therapies for sepsis.

Artesunate (AS) is an effective and reliable anti-malarial drug with low toxicity (Burrows, Chibale & Wells, 2011; Clark, White, S, Gaunt, Winstanley & Ward, 2004), which possesses several interesting effects such as anti-inflammation and other effects (Efferth, Dunstan, Sauerbrey, Miyachi & Chitambar, 2001; Li et al., 2008; Miranda et al., 2013; Wang et al., 2017). Previously, we showed that AS protects septic animals by inhibiting pro-inflammatory cytokine release in cytokine storm stage (Li et al., 2008), suggesting that AS could exhibit anti-inflammatory activity during the cytokine storm phase of sepsis. Herein, cecal ligation and puncture (CLP)-induced sepsis immunosuppression mice model was employed to investigate the effect of AS on sepsis-induced immunosuppression and the lipopolysaccharide/endotoxin (LPS) tolerance cell model was established to unveil the underlying mechanism.

## 2. METHODS

### 2.1 Animals

BALB/c mice (sex: half male and half female; weight: 18 – 22 g; age: 6 – 8 weeks) were obtained from Beijing HFK Bioscience (Beijing, China) and housed in a pathogen-free facility with a 12-h artificial light-dark cycle in the Third Military Medical University. All the mice were provided with food and purified water ad libitum. There were no more than 5 mice in each cage. All animal care and experimental procedures were conducted were approved by Laboratory Animal Welfare and Ethics Committee of the Third Military Medical University (AMUWEC2020015). Animal studies are reported in compliance with the ARRIVE guideline (Kilkenny, Browne, Cuthill, Emerson, Altman & Group, 2010) and with the recommendations made by the British Journal of Pharmacology.

### 2.2 Cell lines, culture and isolation of peritoneal macrophages from mice

The murine macrophage-like cell line RAW264.7 cells and Human monocyte THP-1 cells were purchased from American Type Culture Collection (Manassas, VA, USA). RAW264.7 cells were cultured in Dulbecco’s modified Eagle’s medium (DMEM) supplemented with 10% low- LPS fetal bovine serum (FBS) (Hyclone, Logan, UT, USA), 1% penicillin/streptomycin (regular medium) in a 37°C humid atmosphere with 5% CO_2_. THP-1 cells were cultured in RPMI 1640 (Hyclone) regular medium in suspension. THP-1 derived macrophages were obtained by differentiating THP-1 cells with phorbol 12-myristate 13-acetate (PMA) (Sigma-Aldrich, St. Louis, MO, USA) treatment at 100 nM for 24 h according to the previous study (Li et al., 2017).

Mouse peritoneal macrophages (PMs) were obtained from BALB/c mice. Each mouse was intraperitoneal injected with 3 mL of 3% thioglycolate (Sigma-Aldrich) on day 1 and sacrificed using isoflurane on Day 3. After intraperitoneal injection of 5 mL DMEM regular medium, the peritoneal cells were collected into cell culture dishes. Two hours later, the floating cells were removed by washing the cells with phosphate-buffered saline (PBS). The attached cells were considered to be PMs (purity is about 90%) and were subjected to further experiments. The primary mouse macrophages or cell lines were maintained in regular medium at 37 °C in a humidified 5% CO_2_ atmosphere. All the reagents and utensils used in the experiment were LPS-free.

### 2.3 Bacterial strain and preparation of bacterial suspension

*Pseudomonas aeruginosa* (PA) clinical insolate was kindly provided by Prof. Peiyuan Xia (Southwestern Hospital, Chongqing, China). Single colony was picked from viable, growing Mueller-Hinton (MH) agar plate. Then, it was transferred to 5 ml of liquid MH medium and cultivated aerobically at 37 °C in orbital shaking incubator for 4 h. The cultures were transferred to 100 ml of fresh MH medium for another 12 h. At an OD600 of 0.6 – 0.8, when the bacterial culture was within the logarithmic phase of growth measured by SmartSpec 3000 spectrophotometer (Bio-Rad, Hercules, CA, USA), the suspension was centrifuged at 1500 g for 10 min to harvest the pellet. After washing twice, the pellet was re-suspended by sterile normal saline. The bacteria were diluted in sterile normal saline to achieve the terminal concentration [(colony-formation units (CFU)/mL], which was calculated according to the bacterial growth curve determined by our lab (Concentration = (9.7544 × OD - 1.4965) × 10^8^ CFU/mL).

### 2.4 Establishment of CLP - induced sepsis immunosuppression mice model and AS treatment

Currently, CLP in rodents is regarded as a gold standard in sepsis research (Rittirsch, Huber-Lang, Flierl & Ward, 2009). CLP immunosuppression mice (CLP mice) with second bacterial infection model was established in our lab and described (Deng et al., 2017). Briefly, CLP mice were established by ligating the distal 1.0 cm of the cecum and puncturing once with a No. 16 steel needle, and then were intravenously injected with PA (with 2.5 × 10^9^ CFU/kg) at 24 h after CLP surgery (Figure 1a).

**FIGURE 1.**
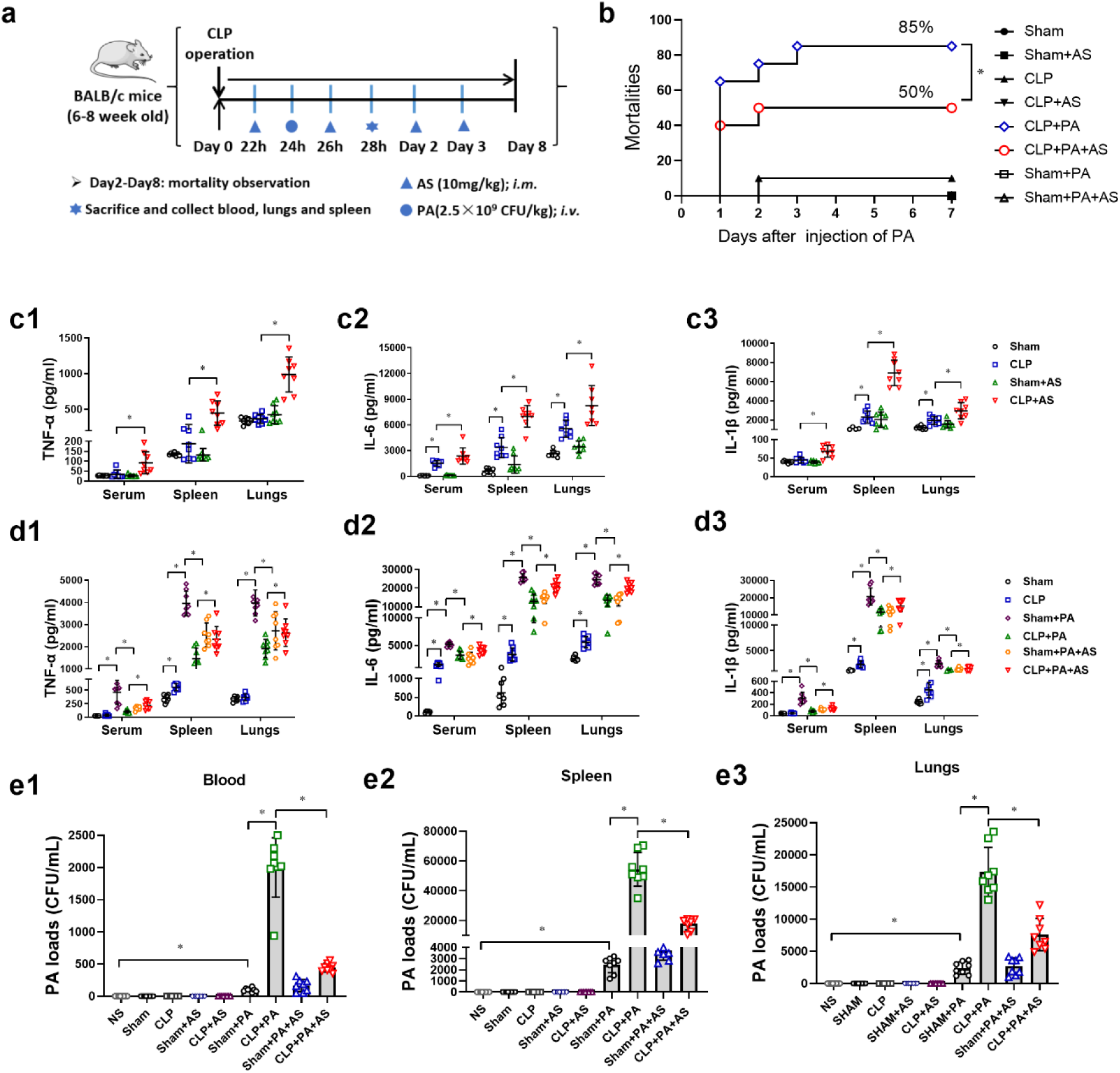
AS reverses sepsis-induced immunosuppression in CLP mice. (a) Schematic diagram to establish CLP mice model with secondary bacterial infection. (b) Mortality of CLP mice challenged with PA and treated by AS (10 mg/kg) (n = 20). (c) Effect of AS treatment on the level of TNF-α (c1), IL-6 (c2), and IL-1β (c3) in the serum, spleen, and lungs of CLP mice (n = 8). (d) Effect of AS treatment on the level of TNF-α (d1), IL-6 (d2), and IL-1β (d3) in the serum, spleen, and lungs of CLP mice challenged with PA (n = 8). (e) Effect of AS treatment on the bacterial load in the blood (e1), spleen (e2), and lungs (e3) 4 h after CLP mice were challenged with PA (n = 8). (* *P* < 0.05; Student’s *t* test)

AS (injection preparation, Guilin Pharma Corp, Guangxi, People’s Republic of China National Medicine Standard H10930195) was added into the attached 5% sodium bicarbonate injection, shaking for 2 minutes until completely dissolved, then diluted in sterile normal saline to achieve the terminal concentration. AS (10 mg/kg) was intramuscularly injected at 22, 26, 48, and 72 h following the CLP surgery. The survival rate of the mice was observed for 7 days. CLP mice were sacrificed at 4 h after PA injection. The blood, lungs, and spleen tissues were collected for CFU count assays and ELISA.

### 2.5 Establishment of LPS-tolerance cell model

LPS tolerance is extensively used to simulate the sepsis-induced immunosuppression phase *in vitro* (Li et al., 2013). Herein, a cell model of LPS tolerance was established in mouse PMs) and RAW264.7 cells. Briefly, cells were cultured with LPS (0111: B4, Sigma-Aldrich, 5 ng/mL) for 4 h. Then the culture supernatant was replaced followed by the addition of LPS (100 ng/mL) to establish the LPS-tolerance cells model.

### 2.6 AS and autophagy inhibitors treatment *in vitro*

For AS treatment, AS (5, 10, and 20 µg/mL) was added with LPS (5 ng/mL) and LPS (100 ng/mL) simultaneously. For autophagy inhibitors treatment, 3-methyladenine (3-MA, 5 mM) (Sigma-Aldrich), bafilomycin A1(Baf, 10 ng/mL) (MedChemExpress, NJ, USA) and LY294002 (Sigma-Aldrich, 10 μM) was added 2 h before the first LPS (5 ng/mL), and also with AS treatment simultaneously. The supernatants and cells were collected at 6 h after the last LPS (100 ng/mL) addition.

### 2.7 siRNA, lentivirus and their transfections *in vitro*

Control siRNA-A (Santa Cruz Biotechnology, Dallas, TX, USA), *VDR* siRNA (Santa Cruz Biotechnology, Dallas, TX, USA) and *ATG16L1* esiRNA (Sigma-Aldrich) were used. RAW264.7 cells were transfected with 50 nM of target siRNA or control siRNA using Lipofectamine 3000 (Thermo Fisher Scientific) and Opti-MEM I reduced serum medium (Thermo Fisher Scientific) for 6 h. After recovering overnight, the cells were subjected to further treatment.

The DNA fragments encoding the mouse VDR (NM_009504) or NF-κB p65 (NM_009045) were synthesized and inserted into the lentivirus vector (#GV358, GeneChem) by GeneChem Corporation (Shanghai, China). The resulting recombinant lentivirus was named VDR-OE or p65-OE. The negative control (#CON238, GeneChem) was named NC-OE. The shRNA targeting mouse VDR (5′-CCTCAAACTCTGATCTGTA-3′), NF-κB p65 (5′-CCCTCAGCACCATCAACTTTG-3′) and the negative control shRNA (5′-TTCTCCGAACGTGTCACGT-3′) were designed, synthesized and inserted into the lentivirus vector (#GV248, GeneChem) by the GeneChem Corporation (Shanghai, China). The resulting recombinant lentivirus was named VDR-KD, NF-κB p65-KD and NC-KD.

For lentivirus transfection, RAW264.7 cells were cultured to 50% confluence and transfected with the lentivirus according to the protocol in the Lentivirus Operation Manual for 6 h. After three days of renewed culture in regular DMEM, the cells were subjected to further treatment.

### 2.8 shRNA transfection *in vivo*

The lentivirus VDR-KD and NC-KD was diluted to a total volume of 200 μl containing 4 × 10^7^ TU. BALB/c mice were transfected through tail intravenous injection as reported (Morizono et al., 2005). One week later, mice were sacrificed and spleen tissues were collect to detect the infection efficiency. Further CLP modeling and AS treatment were employed in transfected mice, and their tissues were collected for the cytokines and western bolting assays.

### 2.9 Interacting molecular prediction for AS

Interacting molecular prediction for AS was performed against 5,311 proteins using the weighted ensemble similarity (WES) algorithm (Zheng et al., 2015) assembled in the traditional Chinese medicine systems pharmacology database and analysis platform (TCMSP) (Ru et al., 2014). WES was developed to determine a drug’s affiliation for a target by evaluation of the overall similarity (ensemble) rather than a single ligand judgment, which allowed us to discovery scaffold hopping ligands. The ligand structural and physicochemical features of AS were calculated using Chemical Development Kit (CDK) (Steinbeck, Han, Kuhn, Horlacher, Luttmann & Willighagen, 2003) and Dragon software and then used to evaluate the ensemble similarity between AS and ligands of candidate targets. Finally, the standardized ensemble similarities of CDK and Dragon were integrated using a Bayesian network to rank the predicted targets.

### 2.10 Transmission electron microscopy

RAW264.7 cells were treated as indicated above and fixed in 2.5% glutaraldehyde at 4°C overnight and post fixed with 2% osmium tetroxide for 1.5 h at room temperature. After fixation, cells were embedded and stained with uranyl acetate/lead citrate. The sections were examined under a transmission electron microscope (JEM-1400PLUS, Japan) at 60 kV.

### 2.11 Immunofluorescence assay

RAW264.7 cells or PMs plated on the coverslips for 2 h and treated as indicated in Method 2.5. The coverslips were fixed in 4% paraformaldehyde for 20 min, and permeabilized in 0.5% Triton X-100 (Solarbio, Beijing, China) which dissolved in PBS for 5 min (20min for nucleoprotein staining). Then the coverslips were blocked for 1 h at 37°C in PBS containing 3% bovine serum albumin (BSA) and 0.1% Triton X-100, and incubated with primary antibodies (displayed in Supplementary Table 1) at 4 °C overnight, followed by secondary antibodies. The coverslips were counterstained with DAPI (Beyotime, Shanghai, China). Confocal images were captured using Zeiss LSM780 confocal microscope (Jena, Germany) with a Plan-Apo Chromat 63 ×/1.40 oil objective.

### 2.12 Methyl thiazolyl tetrazolium (MTT) assay

RAW264.7 cells (5 × 10^4^ cells/well) were plated in 96-well plates in regular medium. After overnight culturing, the cells were washed with PBS and incubated with a series concentration of LPS or certain treatment for 4 or 24 h. Subsequently, 20 μl of the MTT solution (5 mg/ml) was added to the medium in a total volume of 200 μl. The cells were incubated for 4 h at 37 °C in 5% CO_2_. Then, the supernatant was removed, and 150 μl of DMSO was added to each well to dissolve the produced formazan crystals. The extinction was measured at 490 nm using SmartSpec 3000 spectrophotometer (Bio-Rad, Hercules, CA, USA).

### 2.13 Bacterial load assay

For the mouse tissue samples, at the indicated time points after PA injection, the blood, lungs, and spleen were collected. Blood or homogenates of cells, lungs and spleen tissues were serially diluted. Subsequently, 100 μL of the diluents was added to nutrient agar plates and cultured for 18 h. Images of the agar plates and CFU counts were obtained using a G6 automated colony-counter (Shineso Science & Technology, Hangzhou, China).

For cell samples, 2 h after challenged with PA, the cells were collected and washed by PBS three times. The cells were decomposed by 2 min ultrasonication (ultrasonic time 5 s / gap time 10 s cycling, power 100 W) with ice bath, then centrifugated under 4 °C at 12000 rpm for 5 min. The precipitation was resuspended by MH medium and serially diluted.

### 2.14 Binding assay

Microlon ELISA Plates were coated with antibody against VDR (anti-VDR antibody) (Cell Signaling Technology, Boston, MA, USA) overnight at 4 °C, followed by three washes with PBS. The cell lysate containing the total proteins, including the VDR protein, were added into the plates. After incubating at 37 °C for 2 h, the VDR protein was captured by anti-VDR antibody coated on the plates. Subsequently, the plates were washed three times and V_D3_ (Selleck, Houston, TX, USA) was added. After incubating at 37 °C for 2 h, the plates were washed three times and fluorophore-conjugated AS (designed and synthesized by Chongqing Canbipharma Technology, Chongqing, China) was added. After incubating at 37 °C for 2 h, the plates were washed three times and the fluorescence intensity was observed using a Spectra Max i3x Multifunctional enzyme marker (Molecular Devices, USA) with an excitation wavelength of 360 nm and an emission wavelength of 455 nm. AS with fluorophore 12-(7-oxycoumarinyl-ethoxy) dihydro-artemisinin was named AS I, and AS with 12-(-1H-benzo [de] isoquinoline-1, 3(2H)-dione-2-ethoxy) dihydro-artemisinin was named AS II.

### 2.15 Cytokines and NF-κB p65 assays

For the cell samples, the culture supernatants were collected. For blood samples, the serum was collected. For mouse tissue samples, one gram of lungs and spleen tissue was homogenized then the supernatants were collected. The cytokine levels of TNF-α, IL-6, and IL-1β in all of samples were determined using the appropriate ELISA kits (Thermo Fisher Scientific, Waltham, MA, USA), according to the manufactures’ protocols. For NF-κB p65 level in the nucleus, the nucleoproteins were extracted and detected using the Trans-AM NF-κB kits (Active Motif, Carlsbad, CA, USA).

### 2.16 Real-time PCR

After total RNAs were extracted from cells and tissues using the MagMAX total RNA isolation Kit (Thermo Fisher Scientific) and transcribed into cDNAs using Prime Script (Takara, Dalian, China). Using respective primers (displayed in Supplementary Table 2), Real-time PCR was performed (7900HT Fast Real-Time PCR system or ABI 7500 Real-Time PCR system, Applied Biosystems, Darmstadt, Germany). Expression levels were normalized to β-actin. Reactions were performed in duplicate using Tli RNaseH plus and universal PCR master mix (Takara). The relative expression was calculated by the 2(^−ΔΔ**Ct**^) method.

### 2.17 Western blotting

The immune-related procedure used comply with the recommendations made by the British Journal of Pharmacology (Alexander et al., 2018). Cell proteins and mice spleens were extracted using RIPA Lysis Buffer (Beyotime, Shanghai, China). Nuclear and cytoplasmic proteins were isolated by nucleoprotein extraction kits (Keygen Biotech, Beijing, China). Then the proteins were quantified using a BCA kit (Beyotime). Thirty micrograms of each protein sample were separated by 12% SDS-PAGE and transferred to a polyvinylidene difluoride membrane. The membranes were blocked with 5% BSA and incubated with primary antibodies (displayed in Supplementary Table 1) overnight at 4 °C. The membranes were rinsed 5 times with PBS containing 0.1% Tween 20 and incubated for 1 h with the appropriate horseradish peroxidase-conjugated secondary antibody at 37 °C. Membranes were extensively washed with PBS containing 0.1% Tween 20 three times. Chemiluminescence-marked images were developed using the Enhanced Chemiluminescent Substrate (PerkinElmer, Nrowalk, CT, USA) with a ChemiDoc™ Touch Imaging System (Bio-Rad, Hercules, CA, USA) and analyzed using the Image Lab packages (Bio-Rad).

### 2.18 Chromatin immunoprecipitation (ChIP) assays

RAW264.7 cells were washed with cold PBS, crosslinked with 1% formaldehyde and stopped by 0.125 M glycine. Then, cells were sonicated in ChIP lysis buffer to generate 600 bp fragments, followed by an overnight incubation with VDR antibody (Abcam, Cambridge, UK) at 4 °C. Obtained DNA was subjected to semiquantitative or quantitative PCR analysis. Specific primers (displayed in Supplementary Table 3) were used to amplify the VDR-binding region in the promoter of *ATG16L1* genes according to the previous method reported by other laboratories (Wu et al., 2015). ChIP assay was performed using the Pierce Agarose ChIP kit following the manufacturer’s instructions (Thermo Pierce, Rockford, IL, USA) as described before (Bretin et al., 2016).

### 2.19 Coimmunoprecipitation assay

Nuclear and cytoplasmic proteins from RAW264.7 cells were isolated by nucleoprotein extraction kits (Keygen Biotech, Jiangsu, China). Protein (1 mg) was incubated with anti-VDR antibody (Cell Signaling Technology) at 4 °C overnight. Immune complexes were collected using the Protein A/G beads and subjected to SDS-PAGE and western blotting as described above. Coimmunoprecipitation (co-IP) was performed according to the manual of the Protein A/G Magnetic Beads kit (Thermo Fisher Scientific).

### 2.20 Statistical analysis

All the data and statistical analyses comply with the recommendations on experimental design and analysis in pharmacology (Curtis et al., 2018). Briefly, statistical analyses were performed using GraphPad Prism 8.0.2 (263) (GraphPad Software, Inc., CA, USA). All data were expressed as the means ± SD and were analyzed using one-way ANOVA, two-tailed unpaired Student’s *t*-test or Gehan-Breslow-Wilcoxon test. For each parameter of all data presented, * indicates *P* < 0.05.

## 3. Results

### 3.1 AS reverses sepsis-induced immunosuppression in cecal ligation and puncture (CLP) mice

The results from CLP mice showed that AS treatment decreased the mortality of CLP mice challenged with PA from 85% to 50% (Figure 1b), suggesting that AS could protect CLP-induced immunosuppression in mice. In CLP mice unchallenged with PA, the levels of TNF-α, IL-1β, and IL-6 were extremely low in the serum, spleen, and lungs. However, their levels increased significantly in AS-treated CLP mice (Figure 1c). Furthermore, the cytokines levels in CLP mice challenged with PA were also much lower in contrast with sham mice challenged with PA, demonstrating that bacterial challenge could not induce pro-inflammatory cytokines release in sepsis-induced immunosuppressed mice. Importantly, AS treatment significantly augmented the cytokine levels (Figure 1d), indicating that AS could increase cytokine levels in sepsis-induced immunosuppressed mice. Subsequently, the results showed that the bacterial loads obviously increased in the tissues of CLP mice challenged with PA compared with those in the sham mice, demonstrating that the CLP mice had a lower bacterial clearance ability. However, AS significantly decreased the bacterial load (Figure 1e), suggesting AS could protect sepsis-induced immunosuppression by increasing pro-inflammatory cytokine release and decreasing the bacterial load.

### 3.2 AS increases pro-inflammatory cytokines release and bacterial clearance in LPS tolerance macrophages

Based on the concentration-effect relationship between LPS and cytokine production (Figure 2a), 5 ng/mL LPS of four-hour incubation followed by 100 ng/mL LPS was determined to induce LPS tolerance phenotype (tolerance cells) (Figure 2b). Compared with cells treated with only 100 ng/mL LPS treatment (LPS cells), tolerance cells pretreated with 5 ng/mL LPS and followed by 100 ng/mL LPS released much less TNF-α and IL-6 (Figure 2c). However, AS exhibited a dose-dependent enhancement of TNF-α and IL-6 release in tolerance cells (Figure 2d). The results showed the bacterial loads in LPS tolerance cells increased relative to LPS cells at 2 h after PA challenge. Significantly, AS decreased the bacteria load in tolerance cells (Figure 2e). Noteworthily, since the concentrations of LPS (2.5 - 200 ng/mL) and AS(0.625 - 40 μg/ml) used in the experiment had no effect on cell viability, the above effect was not related to the toxicity of LPS and AS on cell activity (Figure S1a) (Li et al., 2008). Subsequent, the human monocyte cell line THP-1 and THP-1 derived macrophages were employed to confirm whether there is a similar effect in human cells. The identical results in the mRNA levels of TNF-α and IL-6 and bacterial clearance in THP-1 cells and THP-1 derived macrophages were obtained (Figure 2f, g). These results were consistent with the findings *in vivo,* demonstrating that AS might reverse sepsis-induced immunosuppression through enhancing pro-inflammatory cytokines release and bacterial clearance by macrophages.

**FIGURE 2.**
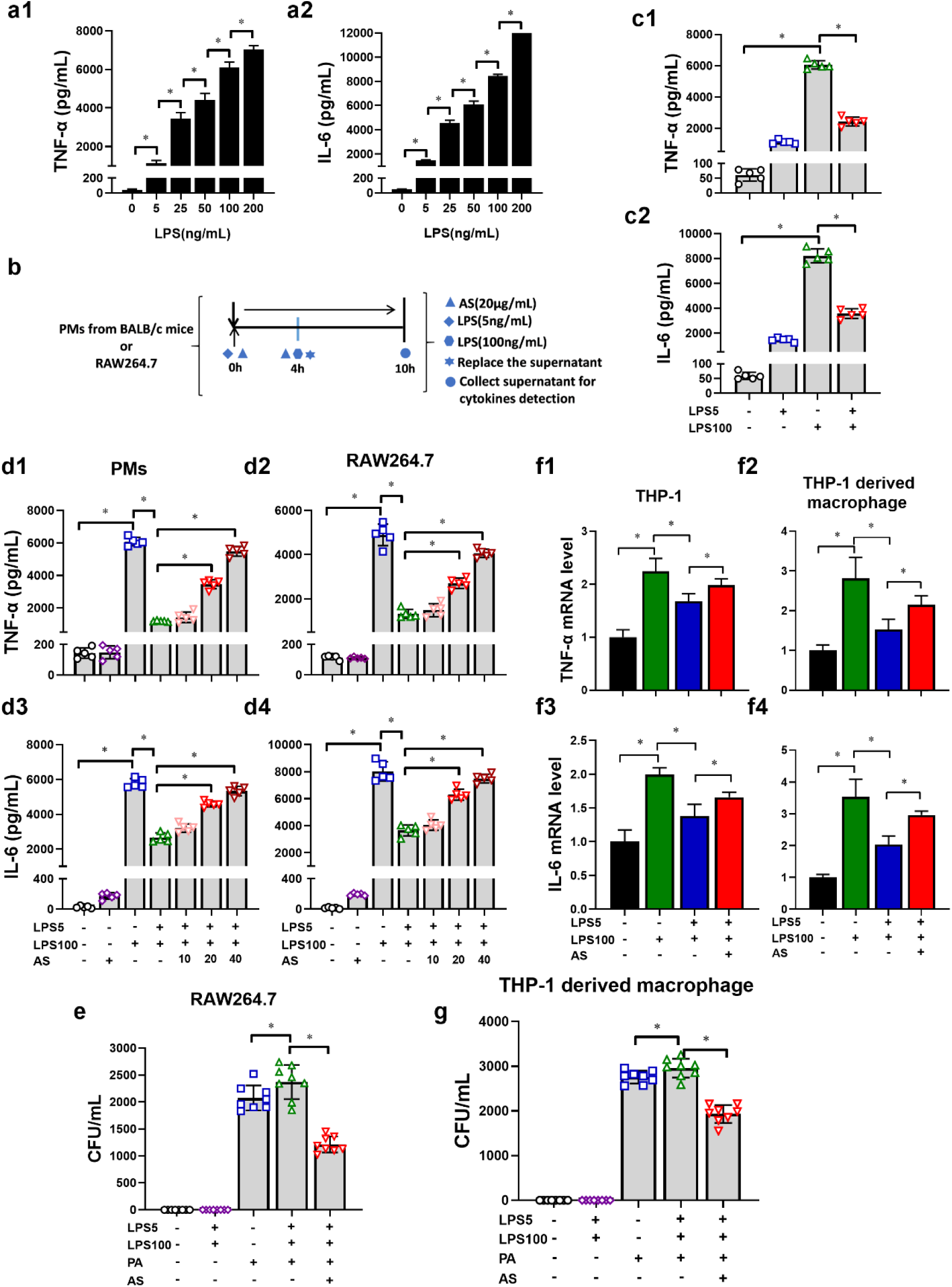
AS increases pro-inflammatory cytokines release and bacterial clearance within LPS-tolerance macrophages (n = 4). (a) LPS increased the release of TNF-α (a1) and IL-6 (a2) from PMs in a dose-dependent manner. (b) Schematic diagram to establish LPS-tolerance macrophages model. (c) The level of TNF-α (c1) and IL-6 (c2) in LPS-tolerance PMs (n = 5). (d) Effect of AS (10, 20 and 40 μg/mL) treatment on the level of TNF-α (d1, 2) and IL-6 (d3, 4) in LPS-tolerance PMs and RAW264.7 cells (n = 5). (e) Effect of AS treatment (20 μg/mL) on the bacterial load in LPS-tolerance RAW264.7 cells (n = 8). (f) Effect of AS (20 μg/mL) treatment on the level of TNF-α (a1, 3) and IL-6 (a2, 4) in LPS-tolerance THP-1 monocytes and THP-1 derived macrophages (n = 5). (g) Effect of AS treatment (20 μg/mL) on the bacterial load in LPS tolerance THP-1 derived macrophages (n = 8). (* *P* <0.05; Student’s *t* test)

### 3.3 VDR is predicted to be an interacted candidate of AS

To further investigate the molecular mechanism of AS, interacted candidates of AS were predicted. Firstly, the top 20 candidates (Figure 3a) were preliminarily screened using RT-PCR. Among them, the most significant and stable change was observed in the mRNA and protein level of VDR, which remarkably increased in tolerance cells and decreased by AS (Figure 3b, c).

**FIGURE 3.**
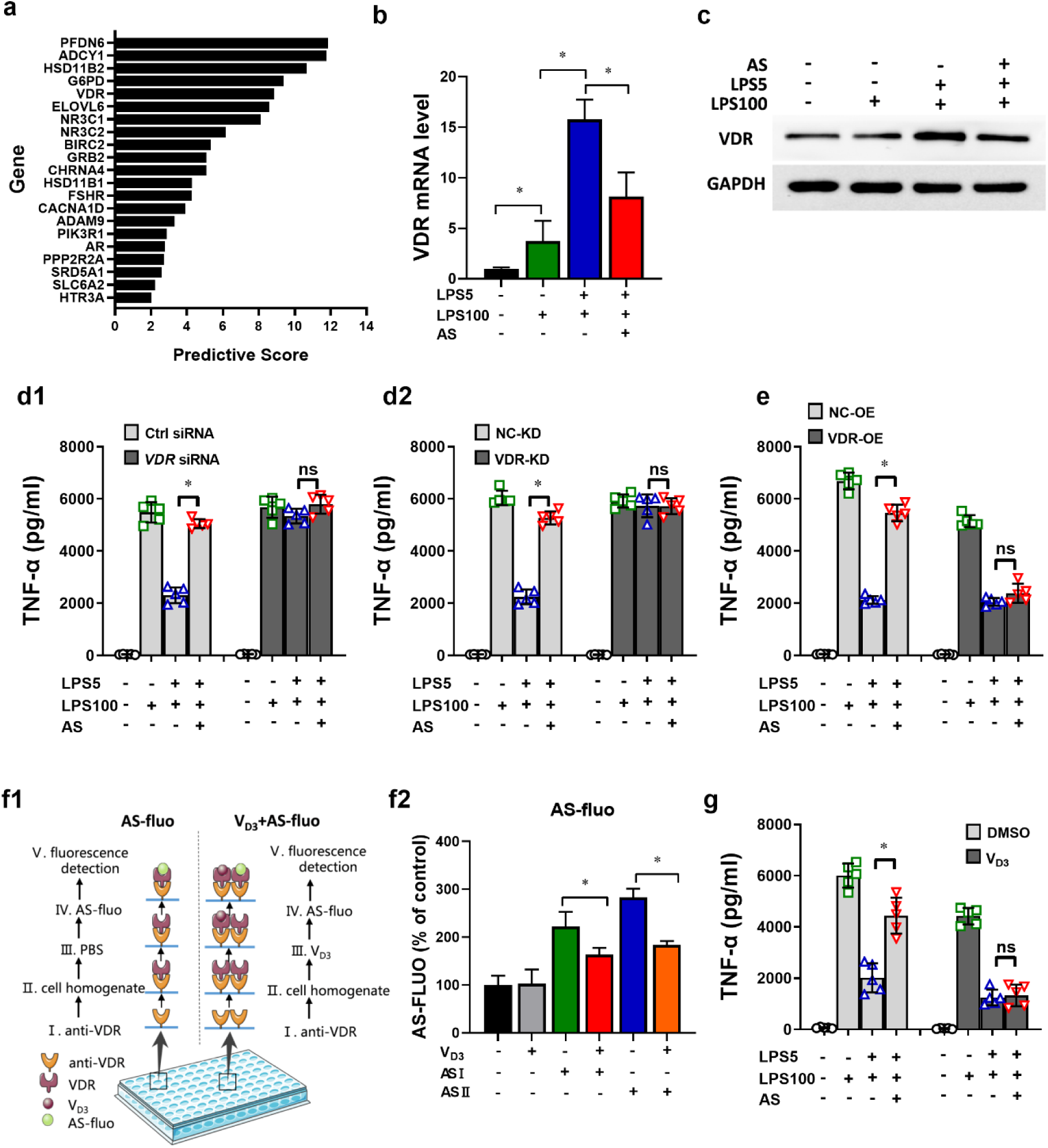
Vitamin D receptor is predicted to be a target candidate of AS. (a) A total of 20 underlying signal molecules were selected via the traditional Chinese medicine systems pharmacology database and analysis platform (TCMSP). (b - c) Effect of AS on the relative level of VDR mRNA (b) and protein (c) expression (n = 5). (d) Effect of *VDR* siRNA (d1) and VDR-KD lentiviral vector (d2) on TNF-α level in LPS tolerance RAW264.7 cells treated with AS (n = 5). (e) Effect of VDR-OE lentiviral vector on TNF-α level in LPS tolerance RAW264.7 cells treated with AS (n = 5). (f1) Schematic diagram of the binding assay designed in our lab. (f2) Effect of V_D3_ on the binding of AS and VDR tracked by AS-fluorophore (n = 5). AS with fluorophore 12-(7-oxycoumarinyl-ethoxy) dihydro-artemisinin was named AS I, and AS with 12-(-1H-benzo [de] isoquinoline-1, 3(2H)-dione-2-ethoxy) dihydro-artemisinin was named AS II. (g) Effect of V_D3_ (100 nM) on TNF-α (level in LPS tolerance RAW264.7 cells treated with AS (n = 5). (* *P* <0.05; n.s., not significant; Student’s *t* test)

Secondly, VDR expression in RAW264.7 was modified. A VDR siRNA and a lentiviral vector harboring an RNAi sequence against VDR were transfected into RAW264.7 cells to knock down VDR expression (VDR-KD cells) (Figure S2a). The results showed that knock-down of VDR would lead to a loss of LPS tolerance phenotype, and addition of AS making no difference (Figure 3d). A lentiviral vector harboring the VDR gene was transfected into RAW264.7 cells to overexpress the VDR (VDR-OE cells) (Figure S2b). The results showed that AS-promoting TNF-α level in LPS tolerance cells was abrogated by overexpressing VDR (Figure 3e). These findings indicated VDR might be an essential and key molecule in LPS tolerance formation and could be further considered as an interacted candidate of AS.

Thirdly, to further verify the interaction between AS and VDR, an indirect competitive binding assay using AS labeled with fluorophore was carried out (Figure 3f1). The results showed pre-incubation of V_D3_ decreased the AS-related fluorescence intensity significantlyFigure 3f2), suggesting that V_D3_ firstly occupied the receptor and then decreased the interaction between AS and VDR.

Lastly, based on V_D3_ is a natural agonist of VDR and there are a very high affinity between them (Haussler, Jurutka, Mizwicki & Norman, 2011), the effect of AS on TNF-α level was investigated in presence and in absence of V_D3_ to validate the above findings. The results showed, in the absence of V_D3_, TNF-α level decreased and AS-promoting TNF-α release in LPS tolerance cells. However, when pro-incubated with V_D3_, the effect that AS-promoting TNF-α release was eliminated (Figure 3g), indicating that there is an antagonism between V_D3_ and AS. These results strongly indicate that AS may bind to the same receptor or even similar sites as V_D3_.

Taken together, all the aforementioned findings demonstrated that AS might interacted with VDR, subsequently inhibiting its biological function of VDR. Therefore, we considered that AS might interact with VDR as an antagonist and then reversed LPS tolerance state.

### 3.4 AS inhibits nuclear translocation of VDR and modulates the transcription of its target gene *ATG16L1*

VDR is a nuclear receptor that mediates most biological functions of V_D3_; it can translocate into the nucleus to regulate the transcription of downstream target genes such as *ATG16L1* via binding to their promotors (Baeke, Gysemans, Korf & Mathieu, 2010), and then regulating the V_D3_-mediated regulation of autophagy and antibacterial effects (Sun, 2016). Therefore, nuclear translocation of VDR and its subsequent effect on target gene *ATG16L1* was investigated firstly. The immunoblotting and immunostaining results showed that VDR protein level dramatically increased in the nucleus of LPS tolerance cells compared with LPS cells, which could be reversed by AS (Figure 4a, b). Secondly, a ChIP assay results showed that AS induced binding of VDR to the promoter of *ATG16L1* as analyzed by semi-quantitative PCR and RT-PCR in LPS tolerance cells (Figure 4c, d). Together, these data showed that AS’s effect is closely related to the transcriptional regulation of VDR on *ATG16L1.* Thirdly, the immunoblotting results indicated that the protein level of ATG16L1 in LPS tolerance cells obviously decreased compared with LPS cells, but AS increased its level (Figure 4e), suggesting VDR negatively regulates the transcription of *ATG16L1* in LPS tolerance cells, which could be prevented by AS. Additionally, our results demonstrated that the protein level of ATG16L1 in LPS tolerance cells markedly increased in VDR-KD group compared with Ctrl-KD group, and the effect of AS was diminished (Figure 4f). However, AS-increasing ATG16L1 level in LPS tolerance cells was abrogated by overexpression of VDR (Figure 4g). All the aforementioned findings indicated AS inhibits the function of VDR via preventing the nuclear translocation of VDR, thus influences the transcription of its target genes *ATG16L1*.

**FIGURE 4.**
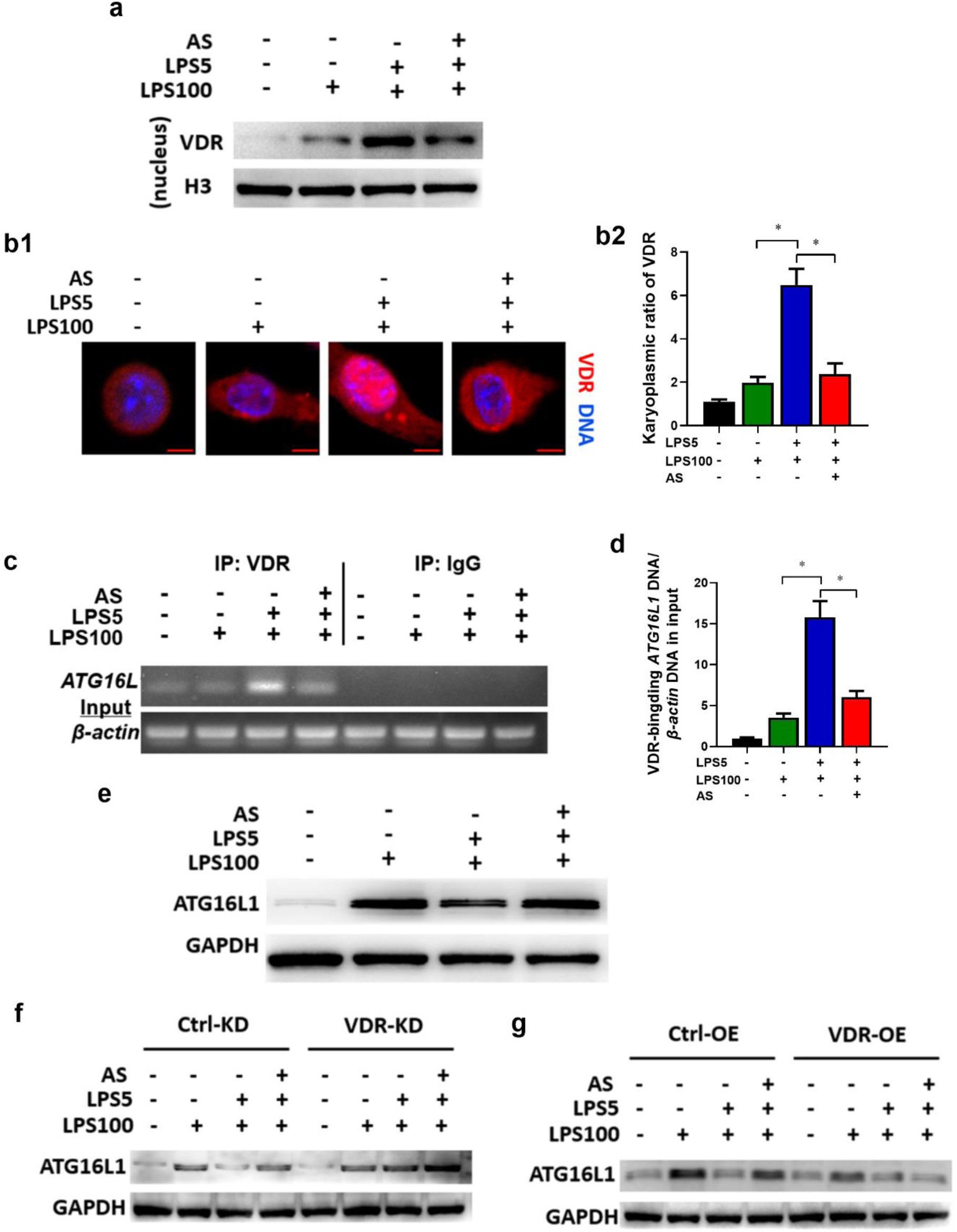
AS inhibits nuclear translocation of VDR and modulates the transcription of its target gene *ATG16L1*. RAW264.7 cells were treated as described in the legend of figure 2d. (a) Immunoblotting to observe VDR level in nucleus lysate. (b) Immunostaining to observe the nuclear translocation of VDR. VDR was probed using Alexa Fluor 555 (red). Representative images (Bar = 5 μm) (b1). The karyoplasmic ratio of VDR was quantified from 100 cells (normalized to medium) (b2). (c) ChIP analysis for the binding of VDR to *Atg16l1* promoter. The protein-DNA complex was immunoprecipitated with anti-VDR or a negative control IgG. Representative agarose gels for the VDR-binding region in *ATG16L1* promoter and *GAPDH* DNA in the input amplified by semiquantitative PCR. (d) The binding of VDR to the Atg16l1 promoter, normalized to Gapdh DNA in the input, analyzed by qPCR (n = 5). (e) The expression of ATG16L1 protein in LPS-tolerance RAW264.7 cells treated with AS. (f) Change of ATG16L1 protein level in LPS tolerance RAW264.7 cells (VDR-KD) treated with AS. (g) Change of ATG16L1 protein level in VDR-OE LPS tolerance RAW264.7 cells treated with AS. (* *P* < 0.05; n.s., not significant; Student’s *t* test)

### 3.5 AS’s effect is autophagy-dependent through VDR *in vitro*

Autophagy is an important self-protection mechanism required for bacterial clearance and inflammation (Gutierrez, Master, Singh, Taylor, Colombo & Deretic, 2004; Levine, Mizushima & Virgin, 2011). ATG16L1, the target gene of VDR and a regulator for autophagy, links VDR and autophagy (Wu & Sun, 2011). Firstly, to confirm the function of ATG16L1, cells were treated with ATG16L1 siRNA (Figure S3a). The results showed that TNF-α and IL-6 levels in LPS cells decreased and AS promoting TNF-α and IL-6 levels in LPS tolerance cells were eliminated in ATG16L1 siRNA infected group relative to negative control (Figure 5a, S2a). It suggests that ATG16L1 is associated with inflammatory cytokine release in LPS tolerance cells and AS’s effects.

**FIGURE 5.**
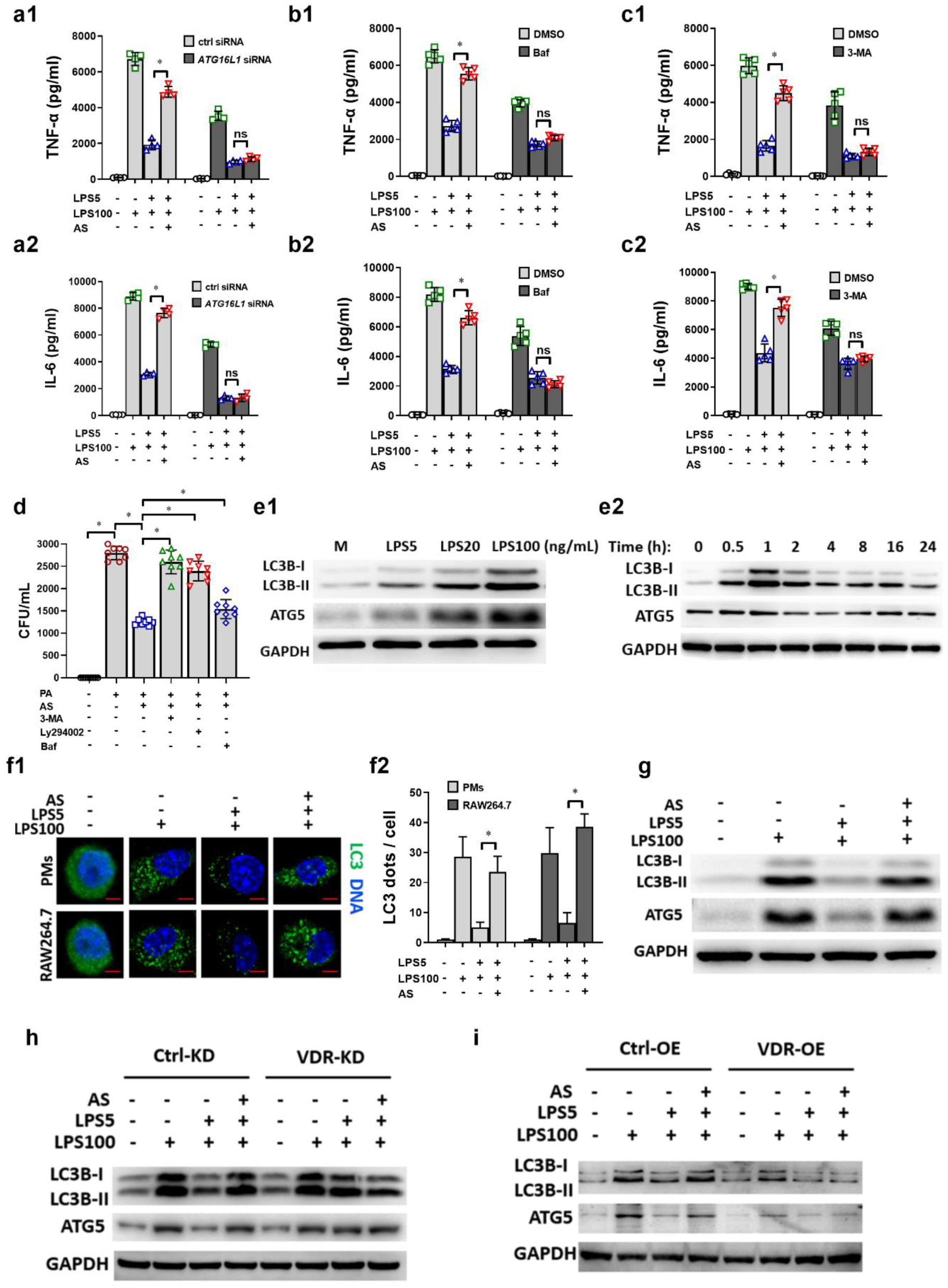
AS’s effect is autophagy-dependent through VDR *in vitro*. (a) Effect of *ATG16L1* siRNA on TNF-α (a1) and IL-6 (a2) level in LPS tolerance RAW264.7 cells treated with AS (n = 5). (b) Effect of Bafilomycin (Baf) (10 ng/mL) on TNF-α (b1) and IL-6 (b1) level in LPS tolerance RAW264.7 cells treated with AS (n = 5). (c) Effect of 3-MA (5 mM) on TNF-α (c1) and IL-6 (c2) levels in LPS tolerance RAW264.7 cells treated with AS (n = 5). (d) Effect of 3-MA, Ly294002 (10 μM) or Baf on the bacteria clearance in LPS tolerance RAW264.7 cells treated with AS (n = 8). (e1) LPS increased the levels of LC3B-I, LC3B-II, ATG16L1 and ATG5 protein expression in a dose-dependent manner in RAW264.7 cells. (e2) The levels of LC3B-I, LC3B-II, ATG16L1 and ATG5 protein expression over time in RAW264.7 cells treated with LPS (100 ng/mL). The level of expression peaked at 1 h. (f) Representative image of immunofluorescence staining of LC3 in LPS tolerance RAW264.7 cells treated with AS (Bar = 2 μm) (f1) Relative fluorescent puncta indicating LC3 aggregation was quantified from 100 cells, the number in the medium group was normalized as 1 (f2). (g) The expression of LC3B-II, ATG16L1, and ATG5 protein in LPS-tolerance RAW264.7 cells treated with AS. (h) Change of LC3B-II, ATG16L1 and ATG5 protein level in LPS tolerance RAW264.7 cells (VDR-KD) treated with AS. (i) Changes of LC3B-II, ATG16L1 and ATG5 protein levels in LPS tolerance RAW264.7 cells (VDR-OE) treated with AS. (* *P* < 0.05; n.s., not significant; Student’s *t* test)

Secondly, three autophagy inhibitors: 3-methyladenine (3-MA) targeting Vps34; bafilomycin A1 (Baf) blocking autophagosome and lysosome fusion; and LY294002 targeting Vps34 (also known as phosphatidylinositol 3-kinase catalytic subunit type 3) were used in RAW264.7 cells. The results showed that TNF-α and IL-6 levels in LPS cells decreased and AS-promoting TNF-α and IL-6 levels in LPS tolerance cells were significantly eliminated when pro-incubated with Baf (Figure 5b) and 3-MA (Figure 5c). Furthermore, our results demonstrated that AS markedly declined the bacteria loads in LPS tolerance cells challenged with PA and this effect could be abrogated when pro-incubated with 3-MA, LY294002 or Baf (Figure 5d). These results indicated a potential correlation between autophagy and AS’s effect.

The above results showed that autophagy contributes to AS’s effect. Thirdly, the influence of AS on autophagy-related molecules was further investigated herein. The markers such as the conversion of microtubule associated protein 1 light chain 3 (LC3B-I) (18 kDa) to LC3B-II (16 kDa) and the accumulation of ATG5 were selected to evaluated autophagy activity (Schaaf, Keulers, Vooijs & Rouschop, 2016). The results showed that LPS treatment increased LC3B-II and ATG5 in a dose-dependent manner (Figure 5e1), consistent with the results that LPS induced pro-inflammatory cytokines release in a dose-dependent manner (Figure 2a). This further indicated that LPS-induced cytokine release is autophagy involved. Immunoblotting showed that accumulation LC3B-II and ATG5 levels changed with time in LPS tolerance cells and peaked at 1 h (Figure 5e2). Therefore, 1 h was selected as the best time point to observe LPS-induced autophagic activities.

Fourthly, immunofluorescence staining, transmission electron microscopy (TEM) and immunoblotting results revealed that autophagy activities remarkably decreased in LPS tolerance cells relative to LPS cells, which could be reversed by AS treatment (Figure S3b, 5f and 5g) at 1 h after the last LPS treatment. These results indicated that autophagy decline is tightly involved in LPS tolerance phenotype and AS treatment potentiated autophagic activities.

Lastly, in LPS tolerance cells, LC3B-II and ATG5 level markedly increased when knocking down VDR compared with negative control, and addition of AS making no difference (Figure 5h). AS-promoting LC3B-II and ATG5 level in LPS tolerance cells was abrogated by overexpressing VDR (Figure 5i). All the aforementioned findings indicated that AS’s effect is autophagy-dependent through VDR *in vitro*.

### 3.6 AS inhibits physical interaction between VDR and NF-κB p65 *in vitro*

NF-κB is an important nuclear transcription factor, and regulates many pro-inflammatory cytokines transcription, both *TNF-α* and *IL-6*, are its target genes (Bonizzi & Karin, 2004). VDR physically interacts with NF-κB p65 in mouse embryonic fibroblast cells and intestinal cells (Sun et al., 2006; Wu et al., 2010), but it remains unclear that the functional relevance of this VDR/NF-κB p65 interaction in LPS tolerance macrophages. Herein, co-IP and fluorescent co-localization experiments were performed. The results showed that the associated NF-κB p65 level in cytoplasm protein had no significant change in the medium cells, LPS cells and LPS tolerance cells when an equal amount of VDR was used as a control (pull-down by VDR), but significantly decreased by AS treatment (Figure 6a1). However, AS had no effect on the nucleus protein of the associated NF-κB p65 in each treatment when an equal amount of VDR was used as a control (pull-down by VDR) (Figure 6a2). These results suggested that AS might bind VDR to interrupt the interaction between VDR and NF-κB p65 in cytoplasm, further implied that AS might affect the activity of NF-κB p65.

Subsequently, immunostaining was used to determine whether AS affected their intracellular co-localization and distribution of VDR and NF-κB p65. The results revealed an obvious co-localization of VDR and NF-κB p65 in LPS tolerance cells, but AS significantly inhibited their co-localization (Figure 6b1, b2). Moreover, nuclear translocation of NF-κB p65 markedly declined in LPS tolerance cells relative to LPS cells, which was reversed by AS treatment (Figure 61, b3). These results indicated that high level of VDR intensively interacted with NF-κB p65, preventing the nuclear translocation of NF-κB p65. Noteworthily, AS inhibited the physical interaction between VDR and NF-κB p65, accelerating the nuclear translocation of NF-κB p65. To confirm it, the nuclear protein level of NF-κB p65 was investigated using ELISA and immunoblotting. The results showed NF-κB p65 level in nucleus obviously reduced in LPS tolerance cells relative to LPS cells, which was reversed by AS (Figure 6c1, c2). In LPS tolerance cells, the nuclear NF-κB p65 level increased in VDR-KD group compared with negative control, leading to the loss of tolerance phenotype (Figure 6d1). In VDR-OE cells, the NF-κB p65 nucleoprotein level remarkably suppressed compared to negative control, and AS treatment made no difference in LPS tolerance cells (Figure 6d2). To determine whether AS-promoting pro-inflammatory cytokines releases in LPS tolerance cells is related to its influence on NF-κB p65 activity, NF-κB p65 expression in RAW264.7 was modified (Figure S4a, b). The results showed that knock-down of NF-κB p65 would lead to the decline in pro-inflammatory cytokines release, and loss of AS’s effect (Figure 6e1). Simultaneously, overexpression of NF-κB p65 would lead to loss of LPS tolerance phenotype, and addition of AS making no difference (Figure 6e2). All the aforementioned findings indicated AS inhibited physically interacts between VDR and NF-κB p65 in LPS tolerance macrophages, and then promoted NF-κB p65 nuclear translocation as well as downstream pro-inflammatory cytokine release.

**FIGURE 6.**
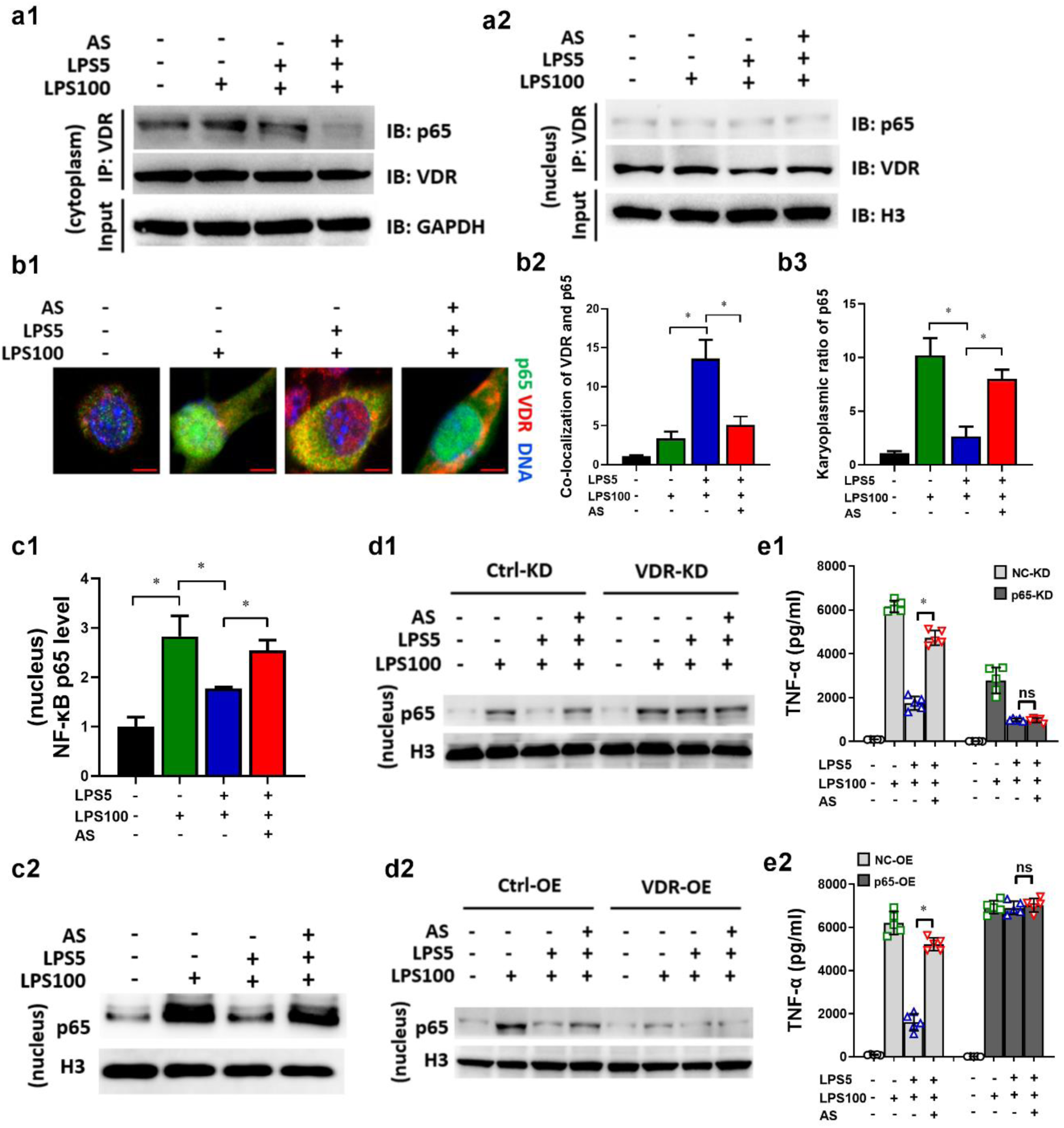
AS inhibits physical interaction between VDR and NF-κB p65 in LPS tolerance macrophages. RAW264.7 cells were treated as described in the legend of figure 2d. (a) The cytoplasm (a1) and nucleus (a2) lysate were used for an IP experiment using anti-VDR antibody, and the associated NF-κB p65 (p65) was detected by immunoblotting (IB). (b) Immunostaining to observe the co-localization of p65 and VDR. P65 was probed using Alexa Fluor 488 (green). VDR was probed using Alexa Fluor 555 (red). Representative images are shown (Bar = 5μm) (b1). The co-localization of VDR and P65 (b2) and karyoplasmic ratio of P65 (b3) was quantified from 100 cells (normalized to medium). (c) P65 level in nucleus lysate were detected using ELISA and IB. (d) Change of P65 level in VDR-KD (d1) or VDR-OE (d2) LPS tolerance RAW264.7 cells treated with AS. (e) Change of TNF-α level in P65-KD (e1) or P65-OE (e2) LPS tolerance RAW264.7 cells treated with AS (n = 5). (* *P* < 0.05; n.s., not significant; Student’s *t* test)

### 3.7 Knock-down of VDR results in the loss of the effect of AS *in vivo*

To correlate the aforementioned findings from *in vitro* experiments, the mRNA and protein levels of VDR were measured in CLP mice. The results showed that the levels of VDR mRNA and protein expression significantly increased in PMs of CLP mice, but were markedly inhibited by AS (Figure 7a), which was consistent with *in vitro* results. In VDR-KD CLP mice which VDR expression was knocked down using a lentiviral vector harboring a VDR RNAi sequence (VDR-KD) (Figure S5a, b), the level of TNF-α in spleen and lungs was significantly increased compare with negative control, and AS administration made no difference (Figure 7b). It suggested that VDR overexpression is involved in sepsis-induced immunosuppression formulation, and knocking down of VDR would lead to loss of immunosuppressing phenotype. Additionally, immunoblotting results showed that autophagy-associated proteins in the spleen of CLP mice were increased compare with negative control, and AS made no difference (Figure 7c). Taken together, both the *in vivo* and *in vitro* results indicate that the VDR plays an important role in sepsis-induced immunosuppression and AS’s effect.

**FIGURE 7.**
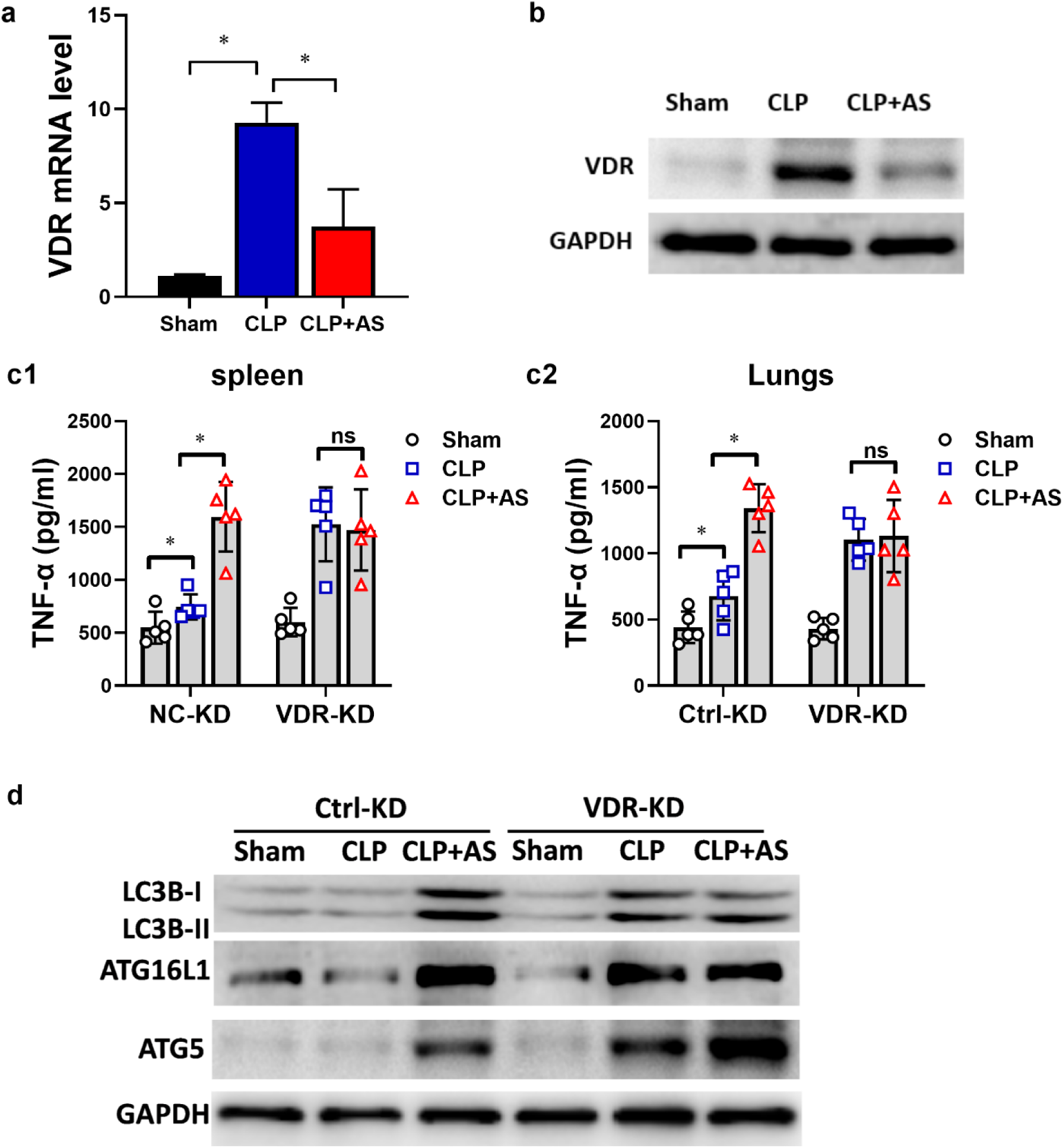
Knockdown of VDR results in the loss of the effect of AS *in vivo*. (a) Effect of AS on VDR mRNA level of PMs from CLP mice (n = 5). (b) Effect of AS on VDR protein level of PMs from CLP mice. (c) Change in the effect of AS on the level of TNF-α and IL-6 in the spleen (c1) and lungs (c2) from VDR-KD mice (n = 5). (d) Changes in the level of LC3B-I, LC3B-II, ATG16L, and ATG5 protein expressions in the spleens of VDR-KD mice. (* *P* <0.05; n.s., not significant; Student’s *t* test)

## 4. Discussion

Herein, our results demonstrated that AS interacted with VDR to reverse sepsis-induced immunosuppression in an autophagy and NF-κB dependent way. Together with our previous results (Kuang et al., 2018; Li et al., 2008), AS is discovered to not only inhibit pro-inflammatory cytokine release during hyper-inflammation stage but also enhance pro-inflammatory cytokines release and bacterial clearance by macrophages to reverse sepsis-induced immunosuppression *in vivo* and *in vitro,* demonstrating that AS exhibits an immunoregulating effect for sepsis treatment, which is not reported previously.

Sepsis is a life-threatening condition caused by a dysregulated host response to infection (Singer et al., 2016). In the last decade, reversing sepsis-induced immunosuppression becomes the focus of sepsis research instead of targeting cytokine storm (Cohen, 2002; Hotchkiss, Monneret & Payen, 2013a; Kox, Volk, Kox & Volk, 2000; Prucha, Zazula & Russwurm, 2017). However, drugs in current clinical trials, such as IL-7, anti-programmed cell death receptor-1 (anti-PD-1) and anti-PD-L1 antibodies, probably only for the immunosuppression stage of sepsis (Hotchkiss, Monneret & Payen, 2013a; Prucha, Zazula & Russwurm, 2017), and its clinical efficacy is yet to be proven.

Clinical investigations have consistently shown a close correlation between Vitamin D deficiency in serum and excess mortality and morbidity, suggesting that vitamin D deficiency is tightly related to sepsis (Dickerson et al., 2016). There are conflicting results from various studies on whether Vitamin D supplementation improves the outcome of patients with sepsis (Kearns, Alvarez, Seidel & Tangpricha, 2015; Putzu et al., 2017). Clinical investigation on the outcome of sepsis patients accepted with Vitamin D supplementation is required in the future. Considering the extensive biological effects of Vitamin D, as well as its role in maintenance of homeostasis in organisms, it is necessary to maintain the physiological concentration of Vitamin D. Based on our results, we consider that low level of Vitamin D might lead to the insufficiency of anti-inflammation response in human body, but excessive Vitamin D might induce the high level of VDR expression; however, the excessive overexpression of VDR might lead to a decreased immune function. Thus, Vitamin D supplementation is necessary in the diet but excessive intake of Vitamin D might be no benefit. Therefore, we consider that an ideal immunotherapy for sepsis is not only to exert an anti-inflammatory effect during the cytokine storm stage, but also increase the resistance to infection during the immunosuppression stage. The therapeutic effects of AS on cytokine storm stage of sepsis and its mechanism has been proved in our previous studies (Kuang et al., 2018; Li et al., 2010; Li et al., 2008). Herein, we revealed that AS possessed therapeutic effects in sepsis-induced immunosuppression *in vivo* and *in vitro*, too.

VDR belongs to a nuclear family receptor. Active VDR binds preferentially as a heterodimer with the retinoid X receptor (RXR) to hexameric repeats on vitamin D response elements (VDRE) in the promoter regions of target genes (Haussler et al., 1998; Orlov, Rochel, Moras & Klaholz, 2012). As with other nuclear receptors, the nuclear translocation of RXR/VDR heterodimer functions to recruit additional cofactors that play an essential role in transcription (Haussler et al., 1998). Herein, VDR was firstly predicted to be an interacted molecule candidate of AS by TCMSP, and then identified using PCR and WB. Moreover, we found that VDR was highly expressed and the nuclear translocation of VDR strikingly increased in LPS tolerance cells (mimic immunosuppression stage of sepsis) compared with LPS cells (mimic cytokines storm stage), while AS treatment could markedly inhibit the high expression and the nuclear translocation of VDR. Furthermore, the interaction between AS and VDR was confirmed in the binding assay established in our lab. These results demonstrated that AS might interact with VDR to affect its activation, leading to a decline of VDR nuclear translocation. Thus, the effects of AS were investigated in presence and in absence of V_D3_, which is the natural ligand of VDR. Our results showed AS’s effect was eliminated when pro-incubated with V_D3_, indicating that there is antagonism between V_D3_ and AS. These results strikingly suggest that AS might bind to the same receptor or even similar sites as V_D3_.

Although the anti-inflammation or immunosuppressive effect of V_D3_ is reported to be tightly related to the function of VDR (Alroy, Towers & Freedman, 1995; Joshi et al., 2011; Lemire, Adams, Kermani-Arab, Bakke, Sakai & Jordan, 1985), few studies have directly addressed the role of VDR in sepsis-induced immunosuppression. Based on our results, we speculated that the excessive high level of VDR during the immunosuppression stage of sepsis might be related to the decreased inflammatory response. While AS could increase the survival rate of sepsis animals via its inhibition on high expression of VDR.

Autophagy is a fundamental cellular process involved in defensive against infection by eliminating intracellular pathogens and regulating inflammation (Casanova, 2017), and autophagy has been considered a new target in sepsis treatment (Ren, Zhang, Wu & Yao, 2017). Herein, combined with our previous study (Kuang et al., 2018), we considered that autophagy was enhanced in the early sepsis stage (cytokine storm stage), but suppressed in the later sepsis stage (immunosuppression stage). Our results also showed AS could enhance autophagy in LPS tolerance cells to eliminate bacteria, and inhibiting autophagy using *ATG16L1* siRNA or autophagy inhibitors could lead to loss of AS’s effects in bacteria clearance and cytokines releases, suggesting autophagy plays a crucial part in AS’s effect.

VDR was reported to be involved in immunoregulation and resistance to infections via regulating autophagy (Baeke, Takiishi, Korf, Gysemans & Mathieu, 2010; Gois, Ferreira, Olenski & Seguro, 2017). VDR transcriptionally regulates *ATG16L1*, which is a target gene and an important autophagy-related protein molecule. If VDR is reduced, ATG16L1 is also reduced associated with abnormal Paneth cells for development of chronic states of mucosal inflammation (Sun, 2016). However, our findings showed that high binding between VDR and *ATG16L1* promotor lead to a low protein level of ATG16L1 in LPS tolerance cells, demonstrating VDR negatively regulated the transcription of *ATG16L1* in LPS tolerance cells. Based on that our results showed AS could inhibit VDR expression to up-regulate the transcription of *ATG16L1* and increase autophagy activity, we considered that the discrepancy for the transcriptional regulation between VDR and *ATG16L1* might be due to the differences in the inflammation and immune status. However, the mechanism needs further investigation in the future.

NF-κB is a vital transcription factor, and the NF-κB signaling pathway links pathogenic signals and cellular danger signals, thus organizing cellular resistance to invading pathogens, which is a network hub responsible for complex biological signaling (Albensi & Mattson, 2000; Kaltschmidt & Kaltschmidt, 2009). NF-κB p65 was reported to physically interact with VDR in mouse embryonic fibroblast cells and intestinal cells (Sun et al., 2006; Wu et al., 2010). Our previous studies have shown that AS could significantly inhibit LPS-induced TLR4/TRA6/NF-κB activation in macrophages, leading to reduced release of proinflammatory cytokines (Kuang et al., 2018; Li et al., 2008), thereby playing an anti-inflammatory role. Herein, we focused on its function in sepsis immunosuppression stage, and discovered that NF-κB p65 was negatively regulated by VDR due to the physical interaction between them in macrophages. Importantly, AS prevented the interaction between VDR and NF-κB p65 in LPS-tolerance cells, leading to a marked increase in nuclear NF-κB p65 translocation and expressions of downstream target genes such as TNF-α and IL-6. Furthermore, we found that AS decreased the binding between VDR and the promoter region of *ATG16L1,* leading to a decline in nuclear translocation of VDR and decreasing the negative transcription of *ATG16L1*, and lastly AS enhanced autophagy activities through binding VDR. Furthermore, knocking down VDR expression reversed the autophagy related proteins levels in immunosuppression stage, and over expressing VDR resulting in a failure of AS-promoting the autophagy related proteins levels. Thus, we concluded that AS interacted with VDR to reverse immunosuppression stage was autophagy involved. Taken together, we think AS interacted with VDR thereby inhibited the nuclear translocation of VDR and also inhibited physical interaction between VDR and NF-κB p65 in LPS tolerance macrophages, then promoted nuclear translocation of NF-κB p65, enhancing autophagy and activating the transcription of NF-κB p65 target genes such as pro-inflammatory cytokines.

As we know, VDR is the receptor to recognize and bind V_D3_. Will V_D3_ control VDR to achieve anti-sepsis effect? Although clinical investigation have shown a close correlation between Vitamin D deficiency in serum and excess morbidity and mortality of sepsis, suggesting vitamin D deficiency is closely related to sepsis (Dickerson et al., 2016), it remains conflicting that Vitamin D supplementation improves the outcome of patients with sepsis (Kearns, Alvarez, Seidel & Tangpricha, 2015; Putzu et al., 2017). Considering the extensive biological effects of Vitamin D, as well as its role in maintenance of homeostasis in organisms, it is necessary and important to maintain the physiological concentration of Vitamin D. Noteworthy, excessive vitamin D supplementation may be harmful rather than beneficial based on our results that excessive Vitamin D might induce the high level of VDR expression, which might lead to decreased immune function.

Till now, the outcome of therapeutic regime only aiming at a single stage of sepsis is disappointing and even worse. It might due to that the treatment time window is important for the therapeutic regimen of sepsis, but difficult to grasp. Giving anti-inflammatory drugs in hypo-inflammatory stage or pro-inflammatory therapy in hyper-inflammatory stage lead to failures even increasing mortalities in clinic (Hotchkiss, Monneret & Payen, 2013a; Rudick, Cornell & Agrawal, 2017). Therefore, we considered that an immunoregulator might be an ideal drug for sepsis without limitation of treatment time window. Our results demonstrated that AS might be an immunoregulator for both hyper- and hypo-inflammatory stage of sepsis based on previous and current results.

Mechanically, AS exerts its “pro-inflammatory” effect in immunosuppression stage in contrast to its “anti-inflammatory” effect in cytokines storm stage. Our present results outline the mechanism of AS’s effect through two ways (presented in Figure 8): AS interacted with VDR to prevent the nuclear translocation of VDR and then decreases its negative regulation of autophagy related target genes such as *ATG16L1*, and then increases autophagy activity. Additionally, AS interacted with VDR to prevent the interaction between VDR and NF-κB p65, and increased the nuclear translocation of NF-κB p65, then augmented the transcription of its target genes such as pro-inflammatory cytokines. Above two ways lead to an increase in the bacteria clearance and the pro-inflammatory cytokine releases of macrophages.

**FIGURE 8.**
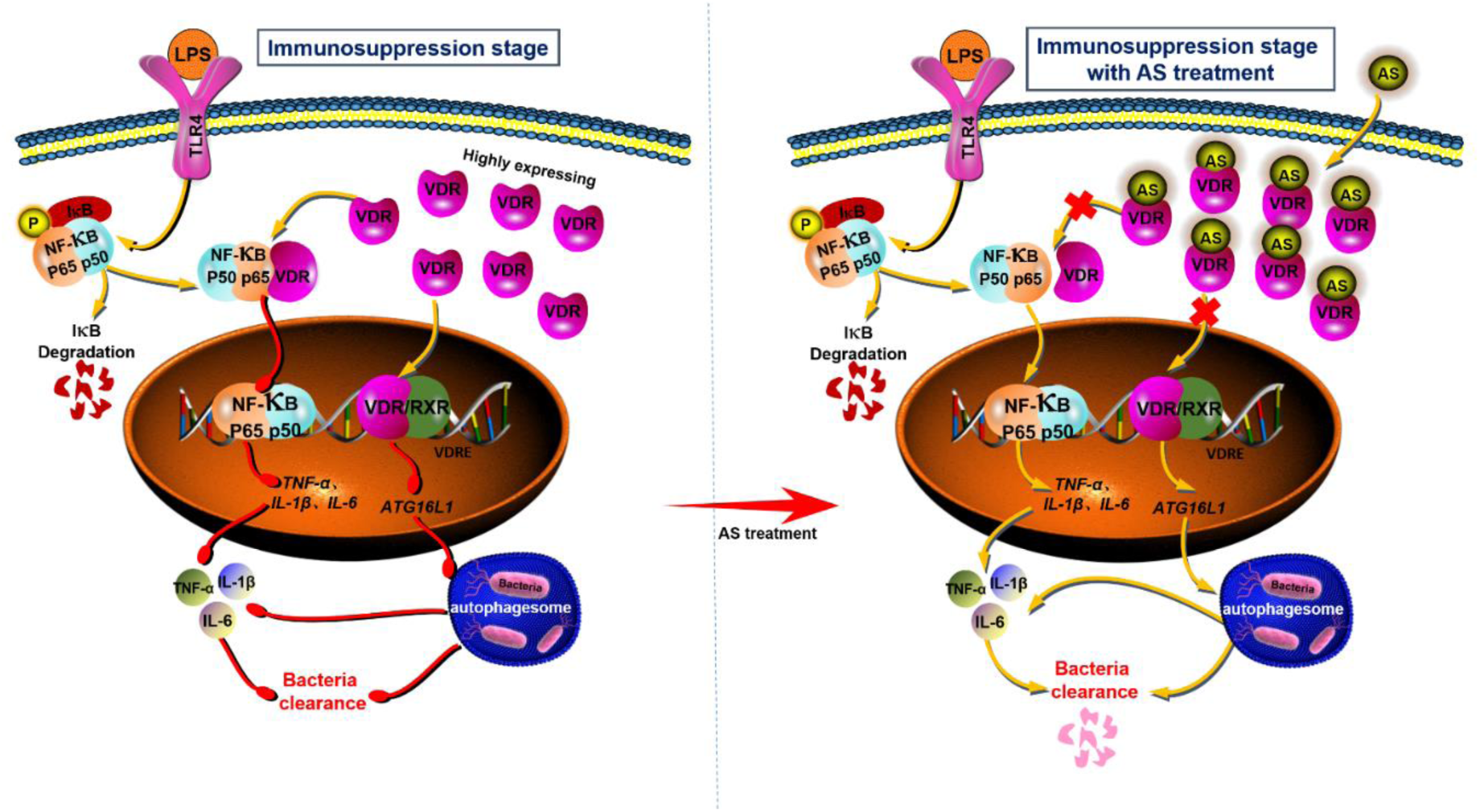
Action mechanism flow chart of AS. AS interacted with VDR to prevent the nuclear translocation of VDR and decrease its negative regulation of autophagy related target genes such as *ATG16L1*, and then increase autophagy activity. Additionally, AS interacted with VDR to prevent the interaction between VDR and NF-κB p65, increase the nuclear translocation of NF-κB p65, then augment the transcription of its target genes such as pro-inflammatory cytokines.

Although monocytes/macrophages system plays a vital role in sepsis, the acquired immune system is equally important, and the effect of AS on T and B cells function remains to be investigated in the future. However, given the therapeutic effect of AS on CLP animal models, we firmly believe that AS will improve the function of T and B cells. VDR is an interaction molecule of AS, whether VDR is a drug target of AS remains to be further studied in the future. Overall, our findings provide an evidence that AS interacted with VDR to reverse sepsis-induced immunosuppression in an autophagy and NF-κB dependent way, highlighting a novel approach for sepsis treatment and drug repurposing of AS in the future.

## Acknowledgements

This work was supported by grants from the National Natural Science Foundation of China (grant number 81872914, 81772137, and 81673495), Major National Science and Technology Program of China for Innovative Drug (2017ZX09101002-002-009), the fourth batch of “Thousand People Innovation and Entrepreneurship Talents Fund” in Guizhou Province, Shijingshan’s Tutor Studio of Pharmacology [GZS-2016(07)]. We sincerely thank Qin Ouyang for his meaningful advice regarding drug targets, Prof. Qiaonan Guo for her electron microscopy assistance and all members of the Zheng laboratory for their help and support. We sincerely thank Dongmei Deng, Bin Li, Mei Kuang, Rongxin Qin, Xichun Pan, Yanyan Cen, Chao Liu, Xiaoli Chen, Zhaoxia Zhai, Yongling Lu and Yan Wang from the Third Military University. We also sincerely thank Huang Chao and Xuetong Chen from the Northwest University for their assistance in this study.

## Author’s contributions

S.S. J.W. and X.L. performed the study under the guidance of H. Z. and S.S. collected and analyzed the data. H. Z. and S.S. wrote the manuscript. X.L., and P.L. contributed to establishing the mouse models. S.S. and L.X. performed the bacteria clearance experiments. S.S. and J.W. were responsible for the molecular biology experiments. Y.W. and X.C. completed the bioinformatics analysis. H.Z., J.Z., S.L. and S.S. contributed to discussions. H.Z., J.Z., and S.L. revised the manuscript. H. Z. is the guarantor of this work, has full access to all the data, and takes full responsibility for the integrity of the data.

## REFERENCES

Albensi BC, & Mattson MP (2000). Evidence for the involvement of TNF and NF-kappaB in hippocampal synaptic plasticity. Synapse 35: 151–159.

Alexander SPH, Roberts RE, Broughton BRS, Sobey CG, George CH, Stanford SC, et al. (2018). Goals and practicalities of immunoblotting and immunohistochemistry: A guide for submission to the British Journal of Pharmacology. Br J Pharmacol 175: 407–411.

Alroy I, Towers TL, & Freedman LP (1995). Transcriptional repression of the interleukin-2 gene by vitamin D3: direct inhibition of NFATp/AP-1 complex formation by a nuclear hormone receptor. Mol Cell Biol 15: 5789–5799.

Angus DC, & van der Poll T (2013). Severe sepsis and septic shock. N Engl J Med 369: 840–851.

Baeke F, Gysemans C, Korf H, & Mathieu C (2010). Vitamin D insufficiency: implications for the immune system. Pediatr Nephrol 25: 1597–1606.

Baeke F, Takiishi T, Korf H, Gysemans C, & Mathieu C (2010). Vitamin D: modulator of the immune system. Curr Opin Pharmacol 10: 482–496.

Bonizzi G, & Karin M (2004). The two NF-kappaB activation pathways and their role in innate and adaptive immunity. Trends Immunol 25: 280–288.

Bretin A, Carriere J, Dalmasso G, Bergougnoux A, B’Chir W, Maurin AC, et al. (2016). Activation of the EIF2AK4-EIF2A/eIF2alpha-ATF4 pathway triggers autophagy response to Crohn disease-associated adherent-invasive Escherichia coli infection. Autophagy 12: 770–783.

Burrows JN, Chibale K, & Wells TN (2011). The state of the art in anti-malarial drug discovery and development. Curr Top Med Chem 11: 1226–1254.

Casanova JE (2017). Bacterial Autophagy: Offense and Defense at the Host-Pathogen Interface. Cell Mol Gastroenterol Hepatol 4: 237–243.

Clark RL, White TE, S AC, Gaunt I, Winstanley P, & Ward SA (2004). Developmental toxicity of artesunate and an artesunate combination in the rat and rabbit. Birth Defects Res B Dev Reprod Toxicol 71: 380–394.

Cohen J (2002). The immunopathogenesis of sepsis. Nature 420: 885–891.

Curtis MJ, Alexander S, Cirino G, Docherty JR, George CH, Giembycz MA, et al. (2018). Experimental design and analysis and their reporting II: updated and simplified guidance for authors and peer reviewers. Br J Pharmacol 175: 987–993.

Deng DM, Li XL, Liu C, Zhai ZX, Li B, Kuang M, et al. (2017). Systematic investigation on the turning point of over-inflammation to immunosuppression in CLP mice model and their characteristics. International Immunopharmacology 42: 49–58.

Dickerson RN, Van Cleve JR, Swanson JM, Maish GO, 3rd, Minard G, Croce MA, et al. (2016). Vitamin D deficiency in critically ill patients with traumatic injuries. Burns Trauma 4: 28.

Efferth T, Dunstan H, Sauerbrey A, Miyachi H, & Chitambar CR (2001). The anti-malarial artesunate is also active against cancer. International journal of oncology 18: 767–773.

Gois PHF, Ferreira D, Olenski S, & Seguro AC (2017). Vitamin D and Infectious Diseases: Simple Bystander or Contributing Factor? Nutrients 9.

Gutierrez MG, Master SS, Singh SB, Taylor GA, Colombo MI, & Deretic V (2004). Autophagy is a defense mechanism inhibiting BCG and Mycobacterium tuberculosis survival in infected macrophages. Cell 119: 753–766.

Haussler MR, Jurutka PW, Mizwicki M, & Norman AW (2011). Vitamin D receptor (VDR)-mediated actions of 1alpha,25(OH)(2)vitamin D(3): genomic and non-genomic mechanisms. Best Pract Res Clin Endocrinol Metab 25: 543–559.

Haussler MR, Whitfield GK, Haussler CA, Hsieh JC, Thompson PD, Selznick SH, et al. (1998). The nuclear vitamin D receptor: biological and molecular regulatory properties revealed. J Bone Miner Res 13: 325–349.

Ho J, Yu J, Wong SH, Zhang L, Liu XD, Wong WT, et al. (2016). Autophagy in sepsis: Degradation into exhaustion? Autophagy 12: 1073–1082.

Hotchkiss RS, Monneret G, & Payen D (2013a). Immunosuppression in sepsis: a novel understanding of the disorder and a new therapeutic approach. Lancet Infect Dis 13: 260–268.

Hotchkiss RS, Monneret G, & Payen D (2013b). Sepsis-induced immunosuppression: from cellular dysfunctions to immunotherapy. Nat Rev Immunol 13: 862–874.

Hutchins NA, Unsinger J, Hotchkiss RS, & Ayala A (2014). The new normal: immunomodulatory agents against sepsis immune suppression. Trends Mol Med 20: 224–233.

Joshi S, Pantalena LC, Liu XK, Gaffen SL, Liu H, Rohowsky-Kochan C, et al. (2011). 1,25-dihydroxyvitamin D(3) ameliorates Th17 autoimmunity via transcriptional modulation of interleukin-17A. Mol Cell Biol 31: 3653–3669.

Kaltschmidt B, & Kaltschmidt C (2009). NF-kappaB in the nervous system. Cold Spring Harb Perspect Biol 1: a001271.

Kearns MD, Alvarez JA, Seidel N, & Tangpricha V (2015). Impact of vitamin D on infectious disease. Am J Med Sci 349: 245–262.

Kilkenny C, Browne W, Cuthill IC, Emerson M, Altman DG, & Group NCRRGW (2010). Animal research: reporting in vivo experiments: the ARRIVE guidelines. Br J Pharmacol 160: 1577–1579.

Kox WJ, Volk T, Kox SN, & Volk HD (2000). Immunomodulatory therapies in sepsis. Intensive care medicine 26 Suppl 1: S124–128.

Kuang M, Cen Y, Qin R, Shang S, Zhai Z, Liu C, et al. (2018). Artesunate Attenuates Pro-Inflammatory Cytokine Release from Macrophages by Inhibiting TLR4-Mediated Autophagic Activation via the TRAF6-Beclin1-PI3KC3 Pathway. Cell Physiol Biochem 47: 475–488.

Lemire JM, Adams JS, Kermani-Arab V, Bakke AC, Sakai R, & Jordan SC (1985). 1,25-Dihydroxyvitamin D3 suppresses human T helper/inducer lymphocyte activity in vitro. J Immunol 134: 3032–3035.

Levine B, Mizushima N, & Virgin HW (2011). Autophagy in immunity and inflammation. Nature 469: 323–335.

Li B, Li J, Pan X, Ding G, Cao H, Jiang W, et al. (2010). Artesunate protects sepsis model mice challenged with Staphylococcus aureus by decreasing TNF-alpha release via inhibition TLR2 and Nod2 mRNA expressions and transcription factor NF-kappaB activation. Int Immunopharmacol 10: 344–350.

Li B, Zhang R, Li J, Zhang L, Ding G, Luo P, et al. (2008). Antimalarial artesunate protects sepsis model mice against heat-killed Escherichia coli challenge by decreasing TLR4, TLR9 mRNA expressions and transcription factor NF-kappa B activation. Int Immunopharmacol 8: 379–389.

Li J, Diao B, Guo S, Huang X, Yang C, Feng Z, et al. (2017). VSIG4 inhibits proinflammatory macrophage activation by reprogramming mitochondrial pyruvate metabolism. Nat Commun 8: 1322.

Li Y, Zhang P, Wang C, Han C, Meng J, Liu X, et al. (2013). Immune responsive gene 1 (IRG1) promotes endotoxin tolerance by increasing A20 expression in macrophages through reactive oxygen species. J Biol Chem 288: 16225–16234.

Miranda AS, Brant F, Rocha NP, Cisalpino D, Rodrigues DH, Souza DG, et al. (2013). Further evidence for an anti-inflammatory role of artesunate in experimental cerebral malaria. Malaria J 12.

Morizono K, Xie Y, Ringpis GE, Johnson M, Nassanian H, Lee B, et al. (2005). Lentiviral vector retargeting to P-glycoprotein on metastatic melanoma through intravenous injection. Nat Med 11: 346–352.

Orlov I, Rochel N, Moras D, & Klaholz BP (2012). Structure of the full human RXR/VDR nuclear receptor heterodimer complex with its DR3 target DNA. EMBO J 31: 291–300.

Prucha M, Zazula R, & Russwurm S (2017). Immunotherapy of Sepsis: Blind Alley or Call for Personalized Assessment? Arch Immunol Ther Exp (Warsz) 65: 37–49.

Putzu A, Belletti A, Cassina T, Clivio S, Monti G, Zangrillo A, et al. (2017). Vitamin D and outcomes in adult critically ill patients. A systematic review and meta-analysis of randomized trials. J Crit Care 38: 109–114.

Ren C, Zhang H, Wu TT, & Yao YM (2017). Autophagy: A Potential Therapeutic Target for Reversing Sepsis-Induced Immunosuppression. Front Immunol 8: 1832.

Rittirsch D, Huber-Lang MS, Flierl MA, & Ward PA (2009). Immunodesign of experimental sepsis by cecal ligation and puncture. Nat Protoc 4: 31–36.

Ru J, Li P, Wang J, Zhou W, Li B, Huang C, et al. (2014). TCMSP: a database of systems pharmacology for drug discovery from herbal medicines. J Cheminform 6: 13.

Rudick CP, Cornell DL, & Agrawal DK (2017). Single versus combined immunoregulatory approach using PD-1 and CTLA-4 modulators in controlling sepsis. Expert Rev Clin Immunol 13: 907–919.

Schaaf MB, Keulers TG, Vooijs MA, & Rouschop KM (2016). LC3/GABARAP family proteins: autophagy-(un)related functions. Faseb J 30: 3961–3978.

Singer M, Deutschman CS, Seymour CW, Shankar-Hari M, Annane D, Bauer M, et al. (2016). The Third International Consensus Definitions for Sepsis and Septic Shock (Sepsis-3). Jama-J Am Med Assoc 315: 801–810.

Skrupky LP, Kerby PW, & Hotchkiss RS (2011). Advances in the management of sepsis and the understanding of key immunologic defects. Anesthesiology 115: 1349–1362.

Steinbeck C, Han Y, Kuhn S, Horlacher O, Luttmann E, & Willighagen E (2003). The Chemistry Development Kit (CDK): an open-source Java library for Chemo- and Bioinformatics. J Chem Inf Comput Sci 43: 493–500.

Sun J (2016). VDR/vitamin D receptor regulates autophagic activity through ATG16L1. Autophagy 12: 1057–1058.

Sun J, Kong J, Duan Y, Szeto FL, Liao A, Madara JL, et al. (2006). Increased NF-kappaB activity in fibroblasts lacking the vitamin D receptor. Am J Physiol Endocrinol Metab 291: E315–322.

Wang J, Zhang J, Shi Y, Xu C, Zhang C, Wong YK, et al. (2017). Mechanistic Investigation of the Specific Anticancer Property of Artemisinin and Its Combination with Aminolevulinic Acid for Enhanced Anticolorectal Cancer Activity. ACS Cent Sci 3: 743–750.

Wu S, Liao AP, Xia Y, Li YC, Li JD, Sartor RB, et al. (2010). Vitamin D receptor negatively regulates bacterial-stimulated NF-kappaB activity in intestine. Am J Pathol 177: 686–697.

Wu S, & Sun J (2011). Vitamin D, vitamin D receptor, and macroautophagy in inflammation and infection. Discov Med 11: 325–335.

Wu S, Zhang YG, Lu R, Xia Y, Zhou D, Petrof EO, et al. (2015). Intestinal epithelial vitamin D receptor deletion leads to defective autophagy in colitis. Gut 64: 1082–1094.

Zheng C, Guo Z, Huang C, Wu Z, Li Y, Chen X, et al. (2015). Large-scale Direct Targeting for Drug Repositioning and Discovery. Sci Rep 5: 11970.

