## supplemental tables and figures for "Artesunate interacts with Vitamin D receptor to reverse mouse model of sepsis-induced immunosuppression via enhancing autophagy"

**Supplementary Figure Legends**

**Figure S1**

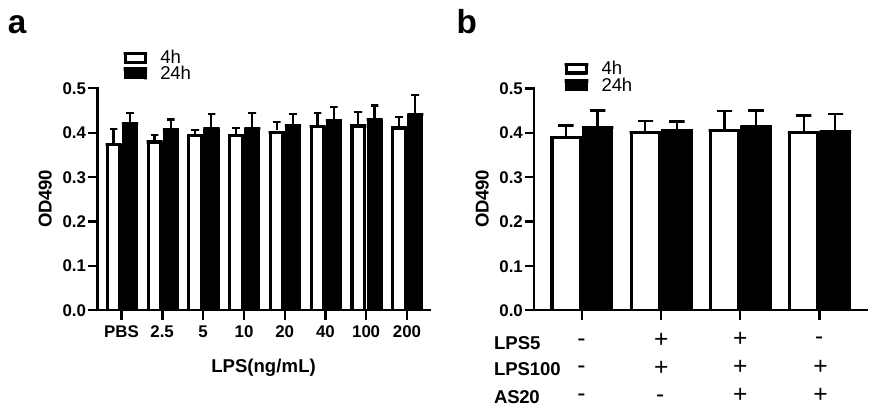
**Figure S1.** Cytotoxicity of LPS on RAW264.7 cells (n=5). (a) the OD 490 value of LPS (2.5 - 200 ng/mL) treatment for 4 h or for 24 h in RAW264.7 cells. (b) the OD 490 value of treatment group (LPS5+LPS100, LPS5+LPS100+AS and LPS100+AS) for 4 h or for 24 h in RAW264.7 cells.

**Figure S2**

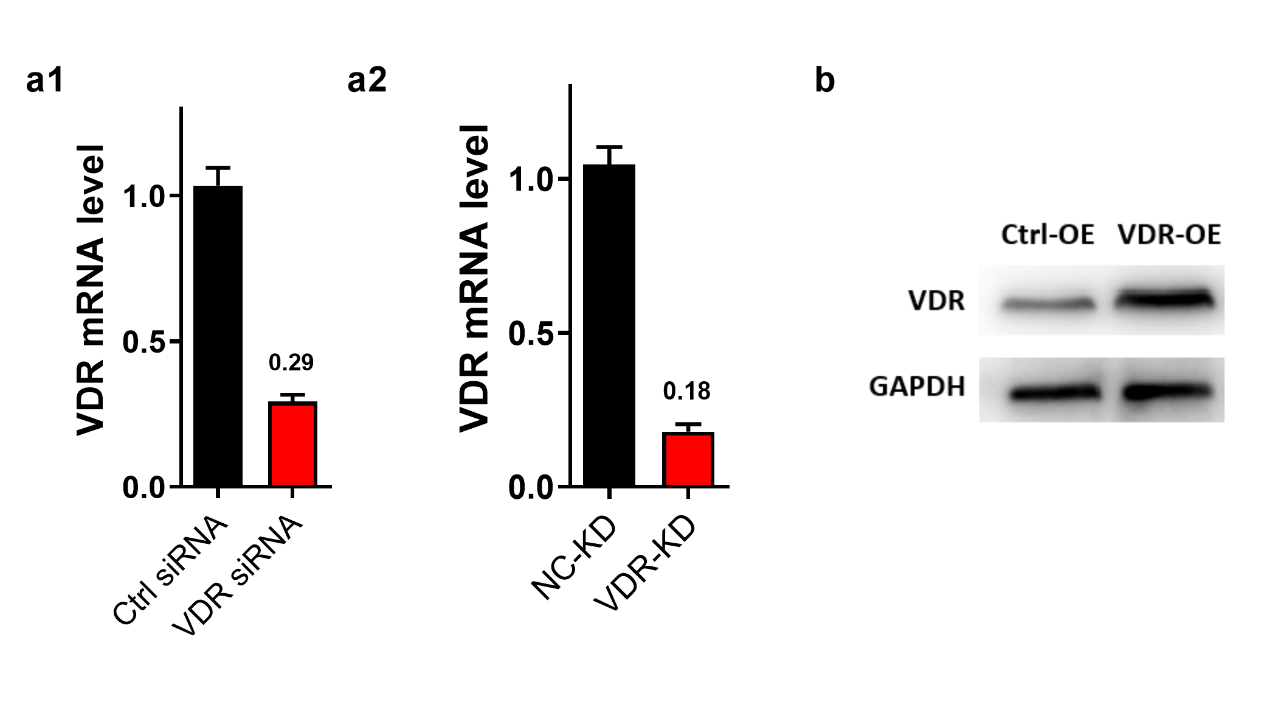

**Figure S2.** Efficiency of modified VDR. (a) VDR mRNA level in the VDR-KD and NC-KD RAW264.7 cells knocked down by VDR siRNA (a1) and lentivirus vector harboring VDR shRNA (a2). (b) Immunoblotting assays of VDR in VDR-OE and Ctrl-OE RAW264.7 cells.

**Figure S3**

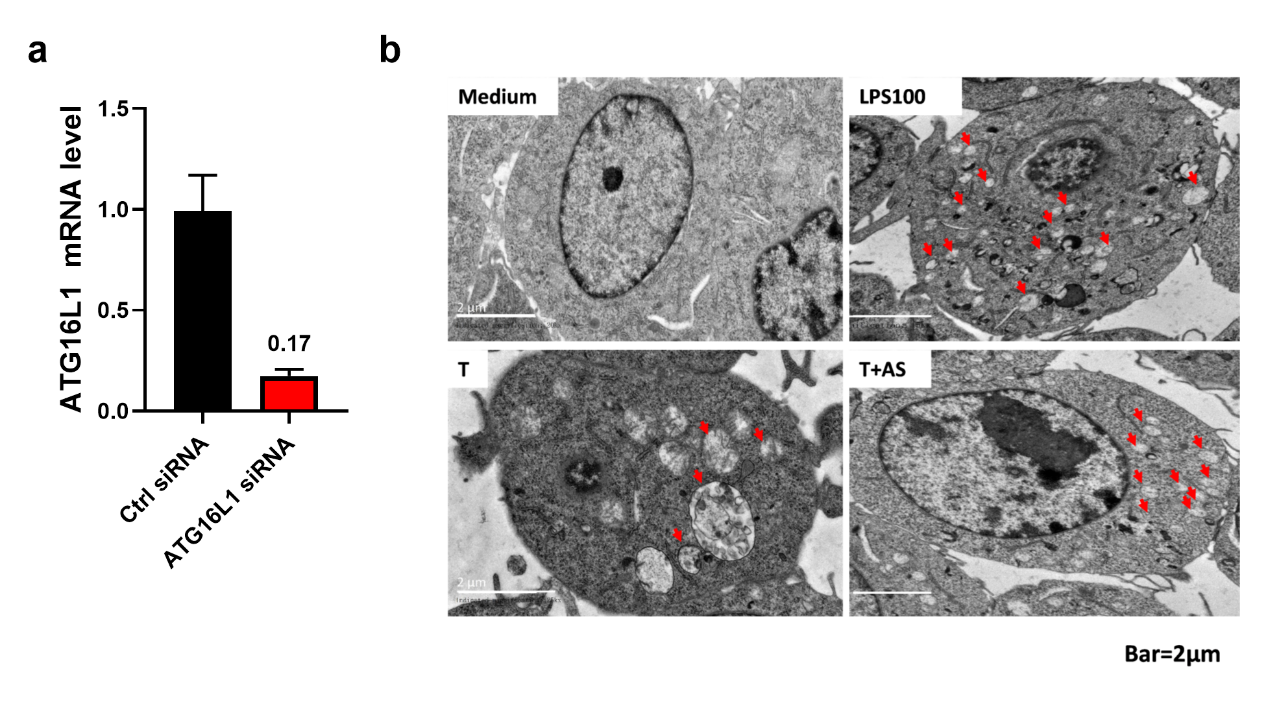

**Figure S3.** AS’s effect is related to autophagy that regulated by VDR. (a) VDR mRNA level in the VDR-KD and NC-KD RAW264.7 cells. (b) Effect of AS on autophagy vesicles in LPS-tolerant RAW264.7 cells under transmission electron microscopy.

**Figure S4**

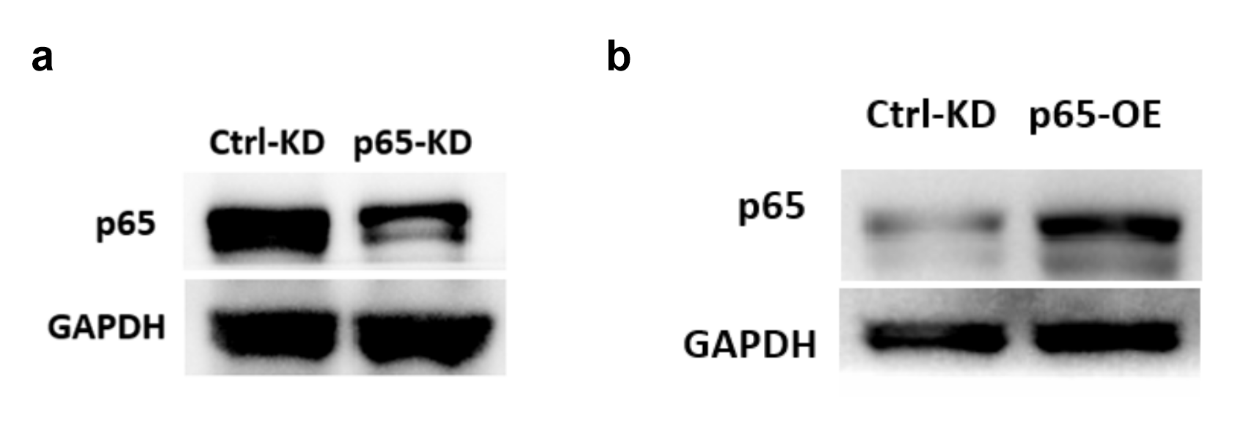

**Figure S4.** Efficiency of modified NF-κB P65. (a) Immunoblotting assays of NF-κB p65 (p65) in P65-KD and NC-KD RAW264.7 cells. (b) Immunoblotting assays of NF-κB p65 in P65-OE and NC-OE RAW264.7 cells.

**Figure S5**

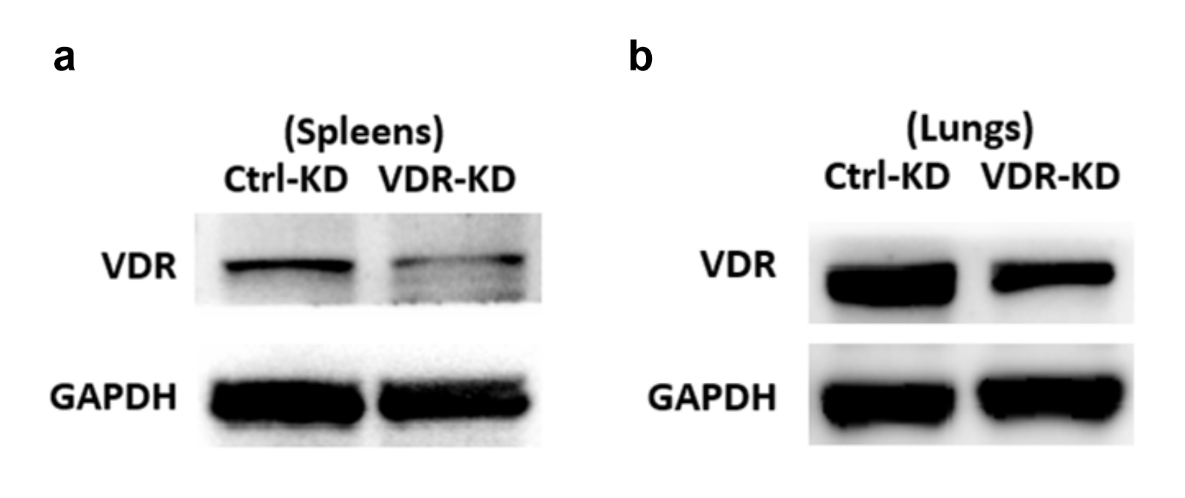

**Figure S5.** Efficiency of VDR knock-down in lungs and spleen. (a) Immunoblotting assays of VDR in the spleen of VDR-KD and NC-KD mice. (b) Immunoblotting assays of VDR in the lungs of VDR-OE and NC-KD mice.

**Supplementary Table 1. Antibodies for Immunofluorescence and Western blotting**

| Antibody | Company | Catalog | Dilution ratio |
| --- | --- | --- | --- |
| anti-NF-κB p65 | Cell Signaling Technology | #9460 | 1:200/1:1000 (IF/WB) |
| anti-VDR | Cell Signaling Technology | #12550S | 1:200/1:1000 (IF/WB) |
| anti-LC3B | Sigma-Aldrich | #L7543 | 1:200/1:5000 (IF/WB) |
| anti-ATG5 | Sigma-Aldrich | #A0731 | 1:1000 (WB) |
| anti-ATG16L1 | Abcam | #ab187671 | 1:1000 (WB) |
| anti-GAPDH | Cell Signaling Technology | #2118 | 1:2000 (WB) |
| goat anti-rabbit IgG (AF 488) | Beyotime | #A0423 | 1:200 (IF) |
| goat anti-mouse IgG (AF 488) | Cell Signaling Technology | #4408 | 1:200 (IF) |
| goat anti-rabbit IgG (AF 555) | Solarbio | #A10704G | 1:200 (IF) |

**Supplementary Table 2. Primer sequences for Real-time PCR**

| Gene | Species | Forward (5' -> 3') | Reverse (5' -> 3') |
| --- | --- | --- | --- |
| *ATG16L1* | mouse | AAGCCGAATCTGGACTGTGG | TATGCAGACTTTGCTGCGGA |
| *VDR* | mouse | GAATGTGCCTCGGATCTGTGG | ATGCGGCAATCTCCATTGAAG |
| *β-actin* | mouse | CGTAAAGACCTCTATGCCAACA | TAGGAGCCAGGGCAGTAATC |
| *TNF-α* | human | TCCTTCAGACACCCTCAACC | AGGCCCCAGTTTGAATTCTT |
| *IL-6* | human | GGCCCTTGCTTTCTCTTCG | ATAATAAAGTTTTGATTATGT |
| *GAPDH* | human | GAAGGTGAAGGTCGGAGTC | GAAGATGGTGATGGGATTTC |

**Supplementary Table 3. promoter specific sequences for *ATG16L1***

| Gene | Species | Forward (5' -> 3') | Reverse (5' -> 3') |
| --- | --- | --- | --- |
| *ATG16L1* | mouse | GGTTCCGTTCTTGTTTCT | TCAAGTTGTCTCCAAGATTAT |
